# CK2 signaling from TOLLIP-dependent perinuclear endosomes is an essential feature of *KRAS* and *NRAS* mutant cancers

**DOI:** 10.1101/2022.04.05.487175

**Authors:** Srikanta Basu, Brian T. Luke, Baktiar Karim, Nancy Martin, Stephen Lockett, Sudipto Das, Thorkell Andresson, Karen Saylor, Serguei Kozlov, Laura Bassel, Dominic Esposito, Mélissa Galloux, Peter F. Johnson

## Abstract

Oncogenic RAS induces perinuclear translocation of the effector kinases ERK and CK2 and their scaffold, KSR1, forming endosomal signaling hubs termed perinuclear signaling centers (PSCs). PSCs are present in all cancer cell lines and tissues examined, suggesting that subcellular compartmentalization of oncogenic kinases drives tumorigenesis. However, the mechanism of perinuclear targeting, whether this location affects kinase substrate specificity, and the importance of PSCs in cancer are unclear. Here we show that the endosomal adaptor, TOLLIP, specifically tethers RAB11A^+^ signaling endosomes containing CK2 and KSR1 to the perinuclear ER. A predicted β-hairpin fold in TOLLIP mediates binding to the KSR1 CA5 pseudo-kinase domain, recruiting CK2/KSR1 complexes to perinuclear endosomes. TOLLIP is essential for proliferation/survival of tumor cells carrying *KRAS* and *NRAS* mutations but not *HRAS*, *BRAF*, *ERBB* or *PTEN* lesions, or non-transformed cells. *KRas^G12D^*-induced lung lesions in *Tollip^-/-^* mice displayed reduced numbers of carcinomatous lesions, implicating TOLLIP in malignant progression. TOLLIP-dependent perinuclear CK2 was shown to phosphorylate discrete substrates, including proteins involved in translation and ribosome biogenesis such as RIOK1. Thus, TOLLIP is a key RAS pathway signaling adaptor in *K/NRAS* tumors whose inhibition is a specific vulnerability of these cancers.

---

*RAS* proto-oncogenes are among the most frequently mutated in cancer, with ∼19% of patients harboring *RAS* alterations^1^. *RAS* cancers are particularly aggressive and treatment-refractory^2,3^. *KRAS* is the most prevalent mutant RAS isoform, representing 75% of all *RAS* cancers. *KRAS* mutations are observed in 88%, 50% and 32% of pancreatic, colon and lung cancer patients, respectively, while *NRAS* alterations are predominant in melanoma and *HRAS* in thyroid and endometrial cancers^1,4^. Oncogenic RAS typically triggers elevated, constitutive signaling through the RAF-MEK-ERK kinase cascade. Studies using integrative pharmacogenomics and phosphoproteomics have also demonstrated an essential role for CK2 kinase in KRAS-driven lung and pancreatic adenocarcinomas (ADCs)^5,6,7^. Despite the importance of RAS effector kinases in tumorigenesis, drugs targeting these kinases have not produced durable therapeutic responses^8,9^. Therefore, it is important to explore additional features of oncogenic RAS signaling that may uncover novel drug targets.

A comprehensive molecular characterization of human lung ADCs showed that p-ERK levels are moderate or low in a substantial proportion of tumor samples, including many that harbor *KRAS* mutations^10^. Similar findings were described for certain lung, colorectal, and pancreatic cancer cell lines^11,12,13,14^ and endometrial tumor samples^15^, suggesting that characteristics besides high MAPK pathway output can contribute to RAS tumorigenesis. Receptors, kinases, scaffolds and RAS proteins can be internalized and signal from endo-membranes^16^. However, the functional consequences of this subcellular compartmentalization, such as whether it influences the substrate selectivity of effector kinases, are unclear. There is also scant information on whether compartmentalized signaling differs in transformed and normal cells. In this regard, we previously reported that RAS-induced transformation coincides with perinuclear re-localization of p-ERK1/2, CK2 and the MAPK scaffold KSR1, which binds to both kinases^17^. These proteins form signaling hubs on nuclear-proximal endosomes termed perinuclear signaling centers (PSCs) that require KSR1 and endosomal trafficking. CK2 and KSR1, but not p-ERK, colocalize with the recycling endosome (RE) marker, RAB11A^17^. In serum-stimulated normal cells, CK2 and p-ERK undergo delayed, transient perinuclear localization, albeit with different kinetics (4 and 6 h post-stimulation, respectively). These findings demonstrate that ERK1/2 and CK2 are associated with distinct types of endosomes.

PSCs are present in all cancer cell lines tested and *KRas^G12D^*-driven mouse lung tumors but not in non-transformed cells or normal lung tissue^17^. These observations suggest that PSC kinases are involved in transducing signals from active RAS, functioning as critical signaling platforms in tumor cells. Therefore, we sought to elucidate the mechanisms that control perinuclear segregation of PSCs, test their functional importance in tumor cells and determine whether PSC-associated kinases selectively phosphorylate specific substrates. We report that the endosomal adaptor and ubiquitin binding protein, TOLLIP, tethers endosomes containing CK2 but not ERK to the perinuclear ER, and that *KRAS* and *NRAS* mutant tumor cells are specifically dependent on TOLLIP for proliferation/survival. We also identify a phosphoproteome requiring perinuclear CK2, demonstrating the importance of subcellular localization in oncogenic signaling.

## Results

### TOLLIP co-localizes with RAB11A^+^ CK2 signaling endosomes in tumor cells

Endocytosis regulates the intracellular sorting, distribution, degradation and recycling of cargo proteins, and modification of endosome trafficking in cancer cells is known to promote tumorigenesis^18,19^. It has been shown that endosomes exist in two spatially distinct but dynamic pools: fast moving peripheral endosomes (PPE) and slow moving perinuclear endosomes (PNE)^20^. PNEs are tethered to the ER through association with the endosomal adaptor proteins EPS15, TOLLIP or TAX1BP1, which bind to mono-ubiquitinated SQSTM1 (p62) on ER membranes. Therefore, we hypothesized that one or more of these adaptors could be involved in perinuclear targeting of signaling endosomes in tumor cells (Supplementary Fig. 1a).

Since KSR1 interacts with multiple RAS effector proteins^21,22,23^ and is required for PSC formation^17^, we asked whether KSR1 binds to endosomal adaptors. Immunoprecipitation experiments in transfected HEK293T cells showed that Pyo-tagged KSR1 associated with endogenous TOLLIP and *vice versa* (Fig. 1a and Supplementary Fig. 1b). This interaction was not stimulated by oncogenic KRAS. Data mining from several published RNAi screens in tumor cell lines revealed TOLLIP as a dependency gene in three studies, two of which involved *KRAS* mutant cells^24,25,26^. These observations prompted us to further investigate the role of TOLLIP in oncogenic signaling.

**Fig. 1.**
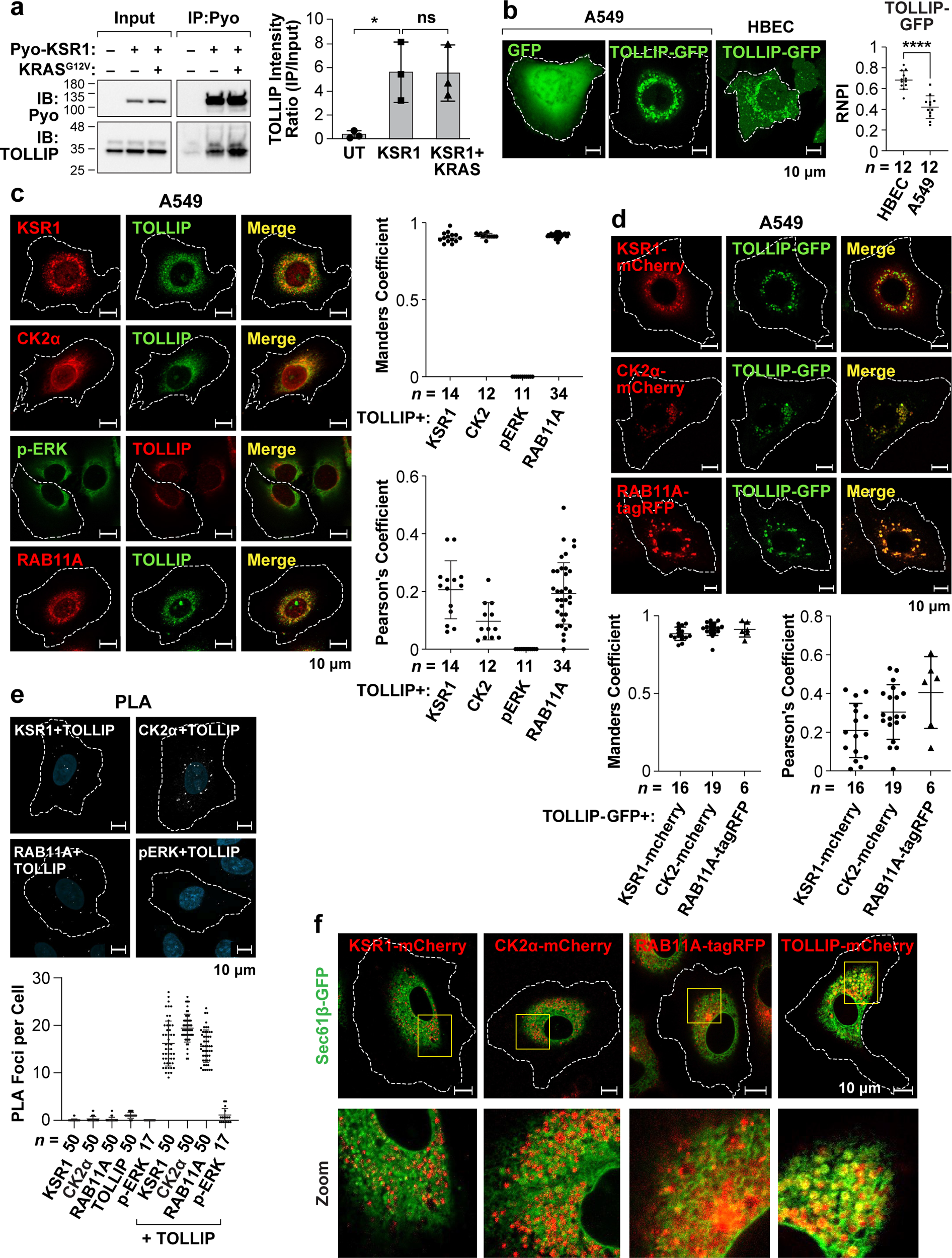
TOLLIP interacts with KSR1 and co-localizes with KSR1 and CK2α on RAB11A+ perinuclear endosomes in tumor cells. **a** Co-immunoprecipitation assay showing association of Pyo-KSR1 with endogenous TOLLIP in 293T cells. Data are mean ± s.d.; *n* = 3 independent experiments. **b** Live cell images demonstrating perinuclear localization of GFP-tagged TOLLIP in A549 lung tumor cells and pan-cytoplasmic distribution in non-transformed bronchial epithelial cells (HBEC). Right: relative nuclear proximity index (RNPI) measures the average distance of fluorescent signals to the nuclear membrane, per cell, normalized to a uniform distribution (see Methods). Data are means for A549 and HBEC; *n*, number of cells analyzed. **c** Immunofluorescence staining (A549 cells) demonstrating co-localization of TOLLIP with KSR1, CK2α and RAB11A but not p-ERK. Right: co-localization of TOLLIP with the other proteins was quantified using Manders *r* and Pearson’s *R*; *n*, number of cells analyzed. **d** TOLLIP-EGFP shows partial co-localization with KSR1-mCherry and nearly complete overlap with CK2α-mCherry and RAB11A-tagRFP in A549 cells. Bottom: co-localization of TOLLIP with KSR1, CK2α and RAB11A was quantified using Manders *r* and Pearson’s *R*; *n*, number of cells analyzed. **e** Proximity ligation assays (PLA) confirm the spatial overlap of TOLLIP with KSR1, CK2α, and KSR1 but not p-ERK in A549 cells. Bottom: PLA signal quantification (foci per cell); *n*, number of cells analyzed. **f** Fluorescently-tagged KSR1, CK2α, TOLLIP and RAB11A are present on endosomal vesicles embedded within the ER network, labeled with Sec61β-GFP, in A549 cells. Statistical significance for co-immunoprecipitation assays and RNPI was calculated using two-tailed unpaired Student’s *t*-test; \**P* ≤ 0.05, \*\*\*\**P* ≤ 0.0001, ns not significant.

Live imaging of EGFP-TOLLIP in A549 cells (human *KRAS^G12S^*lung ADC) showed that TOLLIP localized exclusively to perinuclear puncta, whereas it displayed diffuse, granular pan-cytoplasmic distribution and a perinuclear component in non-transformed HBEC human bronchial epithelial cells (Fig. 1b; Supplementary Videos 1, 2). Nuclear proximity was quantified using an algorithm that measures the distance of cytoplasmic signals from the nuclear envelope, normalized to a uniformly generated distribution for each cell (relative nuclear proximity index, or RNPI). The mean RNPI^TOLLIP^ in A549 cells was 0.4222 and 0.6831 in HBEC cells, indicating a more perinuclear pattern in tumor cells (p<0.0001; Student’s t test). These distinct spatial patterns were also seen for fluorescently-tagged KSR1 and CK2α in the two cell lines (Supplementary Fig. 1c; Supplementary Videos 3, 4), and for endogenous TOLLIP, KSR1 and CK2α (Supplementary Fig. 1d). p-ERK was also perinuclear in A549 cells but its levels were low in HBEC cells, as expected. Furthermore, CK2α, p-ERK and TOLLIP formed juxtanuclear puncta in the pancreatic ductal adenocarcinoma (PDAC) cell line, PANC1 (*KRAS^G12D^*) (Supplementary Fig. 1e).

TOLLIP partially overlapped with KSR1 in A549 cells and displayed prominent co-occurrence with CK2α (Fig. 1c), which was especially evident in live cell imaging (Fig. 1d). However, TOLLIP had minimal overlap with p-ERK. CK2α resides on RAB11A^+^ slow recycling endosomes in tumor cells^17^. Accordingly, IF (Fig. 1c) and live imaging (Fig. 1d) revealed abundant co-localization of TOLLIP with RAB11A, and RAB11A with CK2α and KSR1 (Supplementary Fig. 1f). These conclusions were supported by quantification of signal colocalization using Pearson’s correlation and Manders overlap coefficients. Proximity ligation assays further confirmed the overlap of TOLLIP with CK2α, KSR1 and RAB11A but not p-ERK (Fig. 1e). Taken together, these data show that CK2, TOLLIP, and a fraction of KSR1 localize to perinuclear RAB11A+ endosomes in tumor cells, while ERK resides on a distinct endosomal population.

To test our prediction that PSCs occur on ER-tethered endosomes, we analyzed A549 cells co-expressing TOLLIP-mCherry and the ER marker, EGFP-Sec61β. TOLLIP-positive endosomes were wholly embedded within the perinuclear ER network (Fig. 1f; Supplementary Video 5). Endosomes expressing fluorescently-tagged CK2α, KSR1 and RAB11A were also restricted to this domain. Thus, perinuclear compartmentalization of signaling endosomes involves their intimate association with the ER, which demarcates the PSC region.

We previously observed juxtanuclear localization of KSR1, CK2 and p-ERK in *KRas^G12D^*-driven mouse lung adenomas and ADCs but not in adjacent normal tissue^17^. Similarly, Tollip displayed intense perinuclear staining in lung ADCs but was pan-cytoplasmic in normal lung epithelium (Supplementary Fig. 1g). Tollip was also perinuclear in PDACs arising in *LSL-Kras^G12D^/+*;*p53^R^*^172^*^H^/+*;*Pdx-Cre tg/+* (*KPC*) mice (Supplementary Fig. 1h) and in orthotopic xenografts from a *KPC* PDAC-derived cell line but was diffusely cytoplasmic in normal pancreas (Supplementary Fig. 1i). Thus, the differential subcellular partitioning of TOLLIP seen in tumor vs. non-transformed cell lines extends to *in vivo* tumor tissue.

### TOLLIP is essential for perinuclear targeting of CK2 and tumor cell proliferation

We next asked whether TOLLIP is required for PSC formation in tumor cells. TOLLIP silencing by two different shRNAs in A549 cells disrupted perinuclear localization of KSR1, CK2α and RAB11A, leading to pan-cytoplasmic distribution of each protein (Fig. 2a). However, p-ERK was not significantly affected (Supplementary Fig. 2a). Similar results were seen in PANC1 cells (Supplementary Fig. 2b). TOLLIP silencing also severely reduced the rate of increase in A549 and PANC1 cell populations (Fig. 2b) and another *KRAS* mutant PDAC cell line, MIA PaCa-2 (Supplementary Fig. 2c). The growth defect in A549 cells was associated with increased senescence and apoptosis (Fig. 2c). By contrast, TOLLIP deficiency had little effect on non-transformed HBEC cells (Fig. 2d) or MEFs, as *Tollip^-/-^* cells grew similarly to *WT* controls (Supplementary Fig. 2d).

**Fig. 2.**
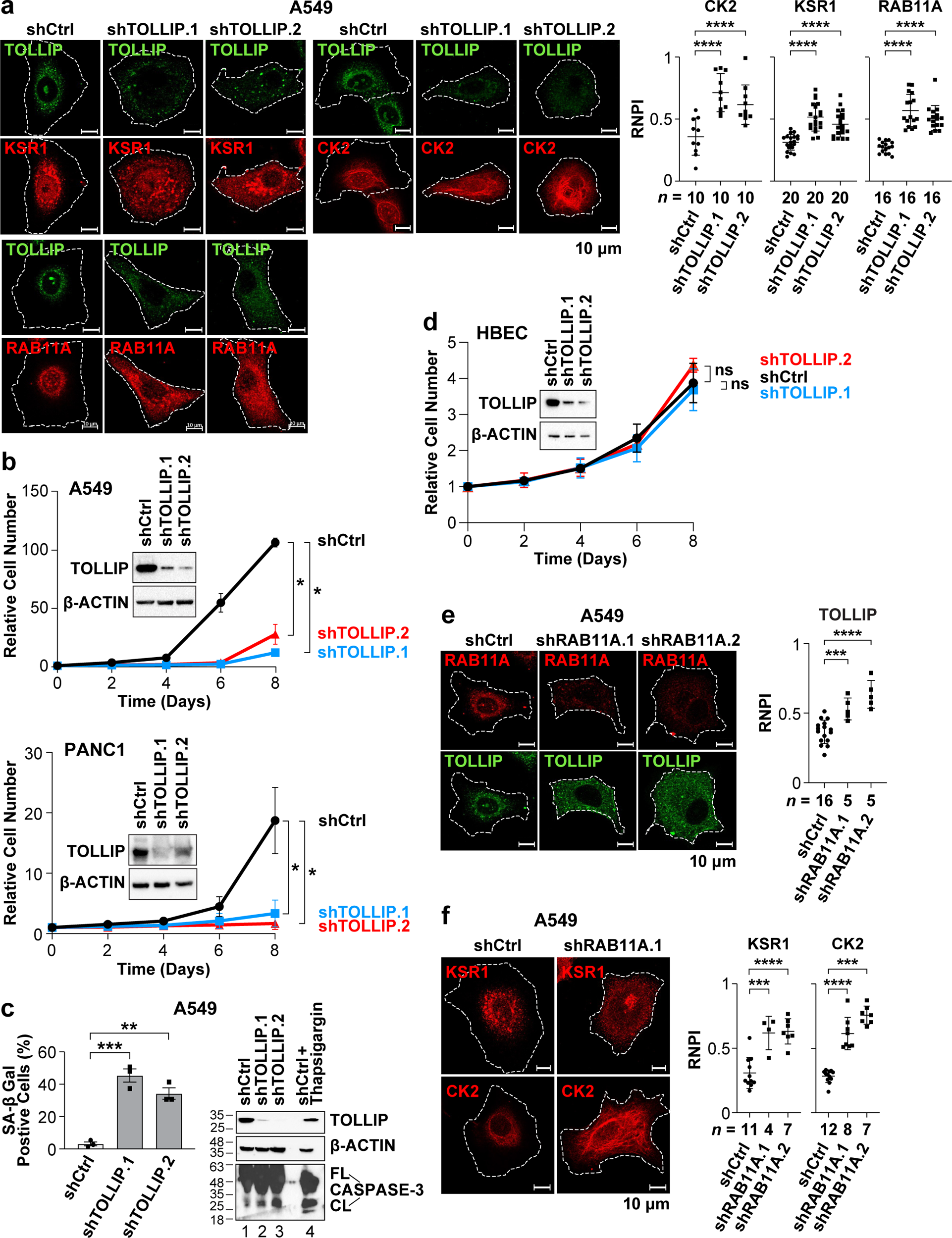
TOLLIP is required for perinuclear localization of CK2α and KSR1 and proliferation/survival of tumor cells but is dispensable in normal cells. **a** TOLLIP depletion in A549 cells causes pan-cytoplasmic dispersal of KSR1, CK2α and RAB11A. Right: RNPI analysis for each protein in control and TOLLIP knockdown cells; *n*, number of cells analyzed. **b** TOLLIP silencing reduces proliferation/survival of A549 and PANC1 tumor cells. Data are mean ± s.d.; *n* = 3 independent experiments. Insets: immunoblots verifying TOLLIP knockdown. **c** TOLLIP-depleted A549 cells display increased senescence (SA-βGal positivity) and apoptosis (cleaved CASPASE-3). SA-βGal data are mean ± s.d.; *n* ≥ 75 cells analyzed per sample. **d** TOLLIP silencing does not affect proliferation of HBEC cells. Data are mean ± s.d.; *n* = 3 independent experiments. Inset shows immunoblots verifying TOLLIP depletion. **e**,**f** RAB11A depletion causes more uniform cytoplasmic distributions of TOLLIP (**e**), KSR1 and CK2α (**f**). A549 cells expressing RAB11A knockdown or control vectors were analyzed by IF staining for the indicated proteins. Right: RNPI analysis for each protein in control and RAB11A-depleted cells; *n*, number of cells analyzed. Two-way ANOVA was used to analyze growth curves. Statistical significance for SA-βGal assays and RNPI was determined using two-tailed unpaired Student’s *t*-test; \**P* ≤ 0.05, \*\**P* ≤ 0.01, \*\*\**P* ≤ 0.001, \*\*\*\**P* ≤ 0.0001.

As TOLLIP, KSR1 and CK2α are associated with RAB11A^+^ endosomes, we investigated whether RAB11A is required for their perinuclear localization. All three proteins became pan-cytoplasmic upon RAB11A silencing (Fig. 2e,f). RAB11A depletion also impaired A549 cell proliferation (Supplementary Fig. 2e). Conversely, TOLLIP was unaffected by KSR1 depletion, although CK2 became dispersed (Supplementary Fig. 2f)^17^. These data suggest that TOLLIP dependency in tumor cells involves its ability, at least in part, to localize CK2 signaling endosomes to the ER. The results support a model in which KSR1 links CK2 to perinuclear RAB11A^+^ endosomes by binding to TOLLIP, which also tethers these vesicles to the ER independently of KSR1.

### A highly conserved sequence in TOLLIP interacts with the KSR1 CA5 domain

Since KSR1-TOLLIP association appears to recruit signaling complexes to perinuclear endosomes, we sought to further characterize this interaction. We generated a nested set of Pyo-tagged KSR1 C-terminal deletion mutants with endpoints at the N-terminal boundaries of five conserved areas (CA1-5) corresponding to known functional domains (Supplementary Fig. 3a)^27^. CA1 binds BRAF and contributes to KSR1 membrane association, CA3 has similarity to atypical C1 lipid binding domains^28^ and binds CK2^23^, CA4 binds ERK^29^, and CA5 is a pseudokinase domain homologous to RAF kinases that interacts with MEK^21,22,30^ and RAF^31^. Co-IP experiments using Pyo-KSR1 deletion mutants expressed with rat HA-Tollip in HEK293T cells showed that removal of CA5 (denoted CA1-4) abolished the KSR1-Tollip interaction (Supplementary Fig. 3a). Interestingly, further deletion of CA4 (CA1-3) partially restored Tollip binding, while loss of CA3 (CA1-2) eliminated the interaction altogether. The CA5 domain alone also bound Tollip efficiently. Thus, KSR1 CA5 appears to be the major site of Tollip interaction. CA3 is also capable of binding Tollip but this association is inhibited by CA4.

To map the corresponding interaction site on Tollip, we performed co-IP assays using Pyo-KSR1 and HA-tagged WT Tollip or various deletion mutants^32^ (Fig. 3a and Supplementary Fig. 3b). TOLLIP contains three known functional domains: TBD (Tom1 binding), C2 (phosphatidylinositol binding) and CUE (ubiquitin binding)^32,33,34,35^ (Fig. 3a). WT Tollip and Δ229-274 (lacking CUE) bound KSR1 efficiently. However, Δ180-274 did not, indicating that the “linker” region between C2 and CUE (residues 179-229) is required for this association. The 179-229 sequence displays marked evolutionary preservation among vertebrates, particularly an 18 aa core (185-202) that is nearly perfectly conserved (Supplementary Fig. 3c). Δ203-274, which adds residues 179-202 to Δ180-274, restored the Tollip:KSR1 interaction while a mutant lacking only residues 180-202 (Δ180-202) lost nearly all KSR1 binding (Fig. 3a). Thus, the invariant sequence between C2 and CUE is essential for KSR1 association. A human TOLLIP structure predicted by AlphaFold2.0^36^ shows a β-hairpin fold corresponding to the conserved 185-202 segment (Fig. 3b). This structure is absent in the model for *C. elegans* Tollip, which lacks this sequence. The precise alignment of the β-hairpin with the 185-202 motif implies that it has a key role in KSR1 binding. A set of Tollip mutants fused to GFP was analyzed for subcellular localization in A549 cells.

**Fig. 3.**
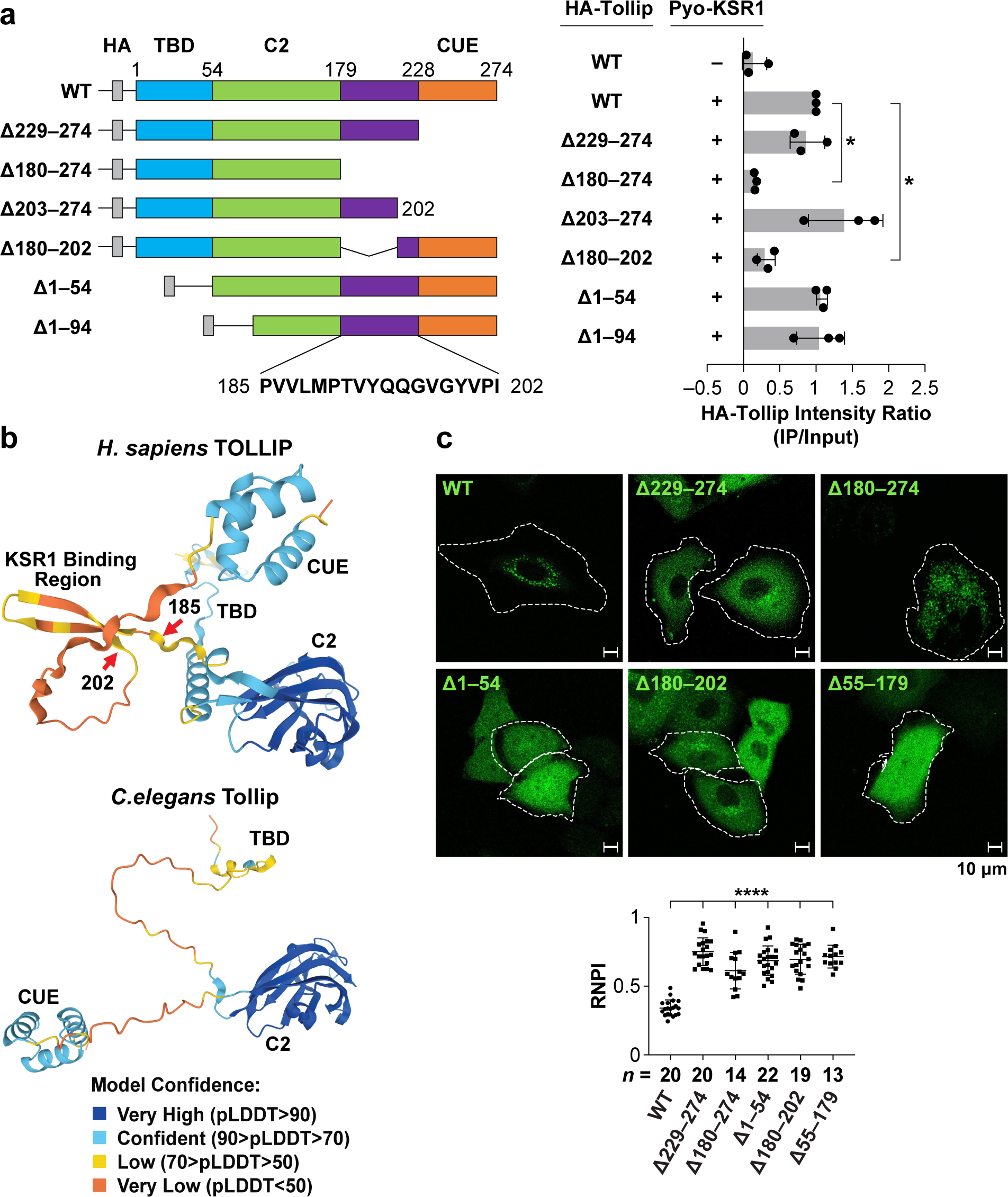
TOLLIP and KSR1 interact through conserved domains in each protein. **a** Mapping the KSR1-interacting region of Tollip. HA-tagged full-length and mutant rat Tollip proteins were co-expressed with Pyo-KSR1 in HEK293T cells. Input lysates and Pyo IP samples were analyzed by immunoblotting for HA (see also Supplementary Fig. 3b) and binding intensity (IP/input) was determined and normalized to WT Tollip. Data are mean ± s.d.; *n* = 3 independent experiments. TBD, TOM1 binding domain; C2, lipid binding domain; CUE, ubiquitin binding domain. **b** A model of human TOLLIP predicted by AlphaFold2^36^ depicts the KSR1 binding region forming a β-hairpin fold. Arrows point to the boundaries of the conserved 18 aa core sequence. *C. elegans* Tollip (bottom), which lacks the conserved KSR1 binding sequence, is not predicted to form the β-hairpin. **c** Subcellular localization of WT Tollip and selected deletion mutants fused to GFP. Proteins were expressed in A549 cells and imaged by fluorescence microscopy. Bottom: RNPI analysis for each protein; *n*, number of cells analyzed. Two-tailed unpaired Student’s *t*-test was performed for IP quantification and RNPI data; \**P* ≤ 0.05, \*\*\*\**P* ≤ 0.0001.

WT Tollip was consistently perinuclear, while each mutant (Δ229-274, Δ180-274, Δ1-54, Δ180-202, and Δ55-179) was more broadly distributed in the cytoplasm (Fig. 3c). All deletions except Δ180-202 impinge on functional domains involved in endosome binding (C2), trafficking (TBD) or ER tethering (CUE) and thus are expected to influence perinuclear localization. Deletion of the KSR1 binding region (Δ180-202) was not anticipated to affect perinuclear targeting. It is possible that resection of the linker region alters folding or accessibility of the flanking C2 and/or CUE domains, thus contributing to the observed localization defect.

### *TOLLIP* is a dependency gene specific to KRAS-driven cancers

As TOLLIP depletion is toxic to *KRAS* mutant tumor cells (Fig. 2), we asked if TOLLIP dependency is specific to *KRAS* cancers or is generally required by transformed cells. Using NIH 3T3 cells transformed by *KRAS^G12V^*or *HRAS^G12V^*, we monitored CK2α and KSR1 localization before and after Tollip knockdown. Both *RAS* oncogenes induced perinuclear translocation of CK2α, KSR1 and Tollip in shCtrl cells (Fig. 4a and Supplementary Fig. 4a). Tollip depletion led to pan-cytoplasmic redistribution of CK2α and KSR1 in NIH 3T3^KRAS^ cells but had little effect in HRAS^G12V^-transformed cells (Fig. 4a). The impact on PSCs correlated with cell proliferation, as Tollip loss decreased the growth rate of NIH 3T3^KRAS^ cells (Supplementary Fig. 4b) but not NIH 3T3^HRAS^ cells (Supplementary Fig. 4c). Proliferation of non-transformed NIH 3T3 cells was unaffected by Tollip silencing (Supplementary Fig. 4b,c). Similar results were seen for *Tollip^−/−^* MEFs, which carried an immortalizing *p19^Arf^*mutation to permit transformation by oncogenic RAS (Supplementary Fig. 4d,e). Growth of *KRAS^G12V^*-expressing *Tollip^−/−^* MEFs was reduced to that of non-transformed cells, whereas *HRAS^G12V^*-tranformed cells were resistant to Tollip deficiency. Consistent with these results, human T24 bladder carcinoma cells (*HRAS^G12V^*) proliferated normally in the absence of TOLLIP and CK2α and KSR1 remained fully perinuclear (Fig. 4b,c). Thus, *KRAS*-driven PSC formation and increased proliferation require TOLLIP, while *HRAS*-transformed cells are TOLLIP independent.

**Fig. 4.**
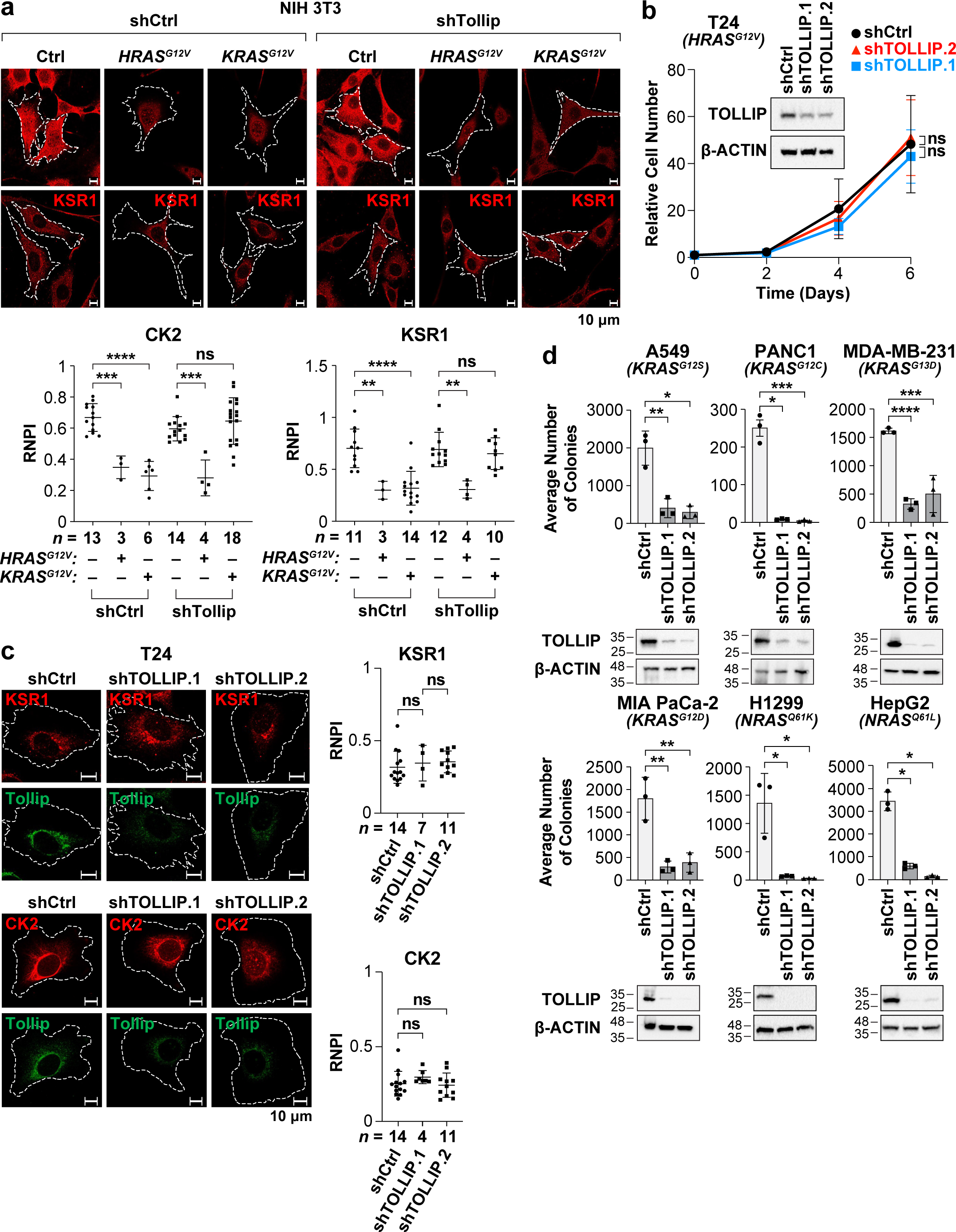
*TOLLIP* is a dependency gene in *KRAS*- and *NRAS*-transformed cells. **a** Control or Tollip-depleted NIH 3T3 cells were transduced with lentiviruses carrying *HRAS^G12V^*, *KRAS^G12V^* or no insert (Ctrl) and immunostained for CK2α and KSR1. Immunoblots confirming Tollip knockdown are shown in Supplementary Fig. 4b,c. Bottom: RNPI analysis for CK2α and KSR1 in each cell population; *n*, number of cells analyzed. **b** TOLLIP silencing does not affect proliferation of T24 cells (human urinary bladder carcinoma; *HRAS^G12V^*). Data are mean ± s.d.; *n* = 3 independent experiments. Inset: immunoblot verifying TOLLIP depletion. **c** Perinuclear partitioning of KSR1 and CK2α in T24 cells is independent of TOLLIP. CK2α and KSR1 localization in control and TOLLIP-depleted cells was analyzed by immunostaining. TOLLIP IF staining confirms silencing efficiency. Right: RNPI analysis of CK2α and KSR1; *n*, number of cells analyzed. **d** Clonogenic growth assays for a panel of human tumor cell lines carrying *KRAS* or *NRAS* mutations, comparing control and TOLLIP knockdown conditions. Data are mean ± s.d., *n* = 3 independent experiments. A549 and H1299, non-small cell lung adenocarcinoma; PANC1 and MIA PaCa-2, pancreatic ductal adenocarcinoma; MDA-MB-231, breast cancer; HepG2, hepatocarcinoma. Oncogenic mutations in each cell line are shown. Immunoblots confirm TOLLIP knockdown efficiency. Growth curves were analyzed by two-way ANOVA. RNPI and clonogenic growth assays were evaluated using two-tailed unpaired Student’s *t*-test; \**P* ≤ 0.05, \*\**P* ≤ 0.01, \*\*\**P* ≤ 0.001, \*\*\*\**P* ≤ 0.0001.

We next compared the effect of TOLLIP depletion across a panel of human tumor cell lines harboring various *RAS* pathway oncogenes. Four *KRAS* and two *NRAS* mutant cell lines showed severely reduced clonogenic growth upon TOLLIP silencing (Fig. 4d). However, RL95-2 endometrial carcinoma (*HRAS^Q61H^*), T24 (*HRAS^G12V^*), A375 melanoma and HT29 colon carcinoma (*BRAF^V^*^600^*^E^*), OVCAR-8 ovarian carcinoma (*HER2^G^*^776^*^V^*) and PC3 prostate cancer (*PTEN/TP53* null) cells were unaffected by TOLLIP knockdown (Supplementary Fig. 5a). These different responses did not correlate with TOLLIP protein levels, which were comparable in the various cell lines (Supplementary Fig. 5b). Thus, tumor cells with *KRAS* or *NRAS* mutations are selectively dependent on TOLLIP. Since CK2 PSCs are induced by oncogenic *HRAS* in NIH 3T3 cells and are also present in human tumor cells carrying oncogenes other than *K/NRAS*, another endosomal adaptor(s) likely provides a redundant ER tethering function in these cells.

### Characterization of a TOLLIP-dependent transcriptome in tumor cells

To further define TOLLIP-regulated cellular processes, we used RNA-seq to compare the transcriptomes of control and TOLLIP-depleted A549 cells (Supplementary Fig. 5c; Supplementary Table 1). Genes that decreased upon TOLLIP knockdown were highly correlated with gene ontology (GO) processes for cell cycle, DNA replication and chromosome segregation (Supplementary Fig. 5d; Supplementary Table 2). Related categories were also identified through gene set enrichment analysis (GSEA; suppressed processes) (Supplementary Fig. 5e; Supplementary Table 3) and include several genes encoding subunits of the MCM helicase complex regulating initiation of DNA replication. This effect may be related to the identification of MCM2 as a potential TOLLIP-dependent CK2 substrate as well as decreased phosphorylation of cyclin-dependent kinase (CDK) targets in TOLLIP-depleted cells (see below). Specifically, reduced MCM2 phosphorylation/activity^37,38^ and diminished CDK signaling could impair the onset of DNA replication and subsequent expression of S-phase MCM genes. Predominant upregulated processes were inflammatory response and immune response (Supplementary Fig. 5f and Supplementary Table 3), possibly reflecting cellular stress and senescence elicited by TOLLIP depletion and consistent with TOLLIP’s role in regulating pro-inflammatory signaling^39^.

### Tollip facilitates progression to adenocarcinoma in a *Kras^G12D^*-driven model of lung cancer

To investigate whether Tollip deficiency impairs *Kras*-induced tumor development *in vivo*, we produced *Tollip^−/−^* mice^41^ carrying a Cre-activated *LSL-Kras^G12D^*oncogene^42^. Ad.Cre virus was administered to the lungs of *WT* or *Tollip* null *LSL-Kras^G12D^*mice and animals were maintained until disease endpoint. All mice eventually succumbed to respiratory distress due to accumulated pulmonary lesions, and no survival difference was observed between genotypes (P=0.40; Log-rank test) (Fig. 5a). Lung lesions in this model are primarily benign alveolar hyperplasia and adenomas, a few of which advance to malignant ADC (Fig. 5b). Hyperplastic areas (Fig. 5c) and adenoma counts (Fig. 5d) were similar between genotypes, with high adenoma burdens accounting for the similar mortality rates between the two strains. However, carcinomatous lesions were significantly reduced in *Tollip^−/−^* mice (Fig. 5d). ADCs were the highest tumor grade observed in 83.33% of *Tollip^+/+^* mice, whereas only 42.1% of *Tollip^−/−^*animals presented with carcinomas (P=0.017; Fisher’s exact test) (Fig. 5e). These findings reveal a significant block to *Kras*-driven ADC development in mutant animals, indicating an important role for Tollip in malignant progression. A previous study reported that Tollip deficiency caused a reduction in colitis-associated colon cancers, which are primarily adenomas^40^. This was attributed to diminished inflammation and reduced infiltration of CD4+/Foxp3+ regulatory T cells and other immune cells. Using CD45 as a pan-immune marker, we did not observe differences in overall immune cell infiltration between *WT* and *Tollip^−/−^* lung adenomas or ADCs (Fig. 5f), suggesting that the cancer phenotype involves tumor-cell-intrinsic Tollip deficiency. To further address the tumor-cell-autonomous role of Tollip, we delivered TOLLIP-depleted A549 cells intravenously into immunodeficient mice. Whereas cells expressing a control shRNA formed an average of 80 lung lesions per animal, two different shTOLLIP constructs almost completely prevented tumor formation and most animals presented no detectable lesions (Fig. 5g). Thus, human lung ADCs are highly dependent on TOLLIP for tumor engraftment.

**Fig. 5.**
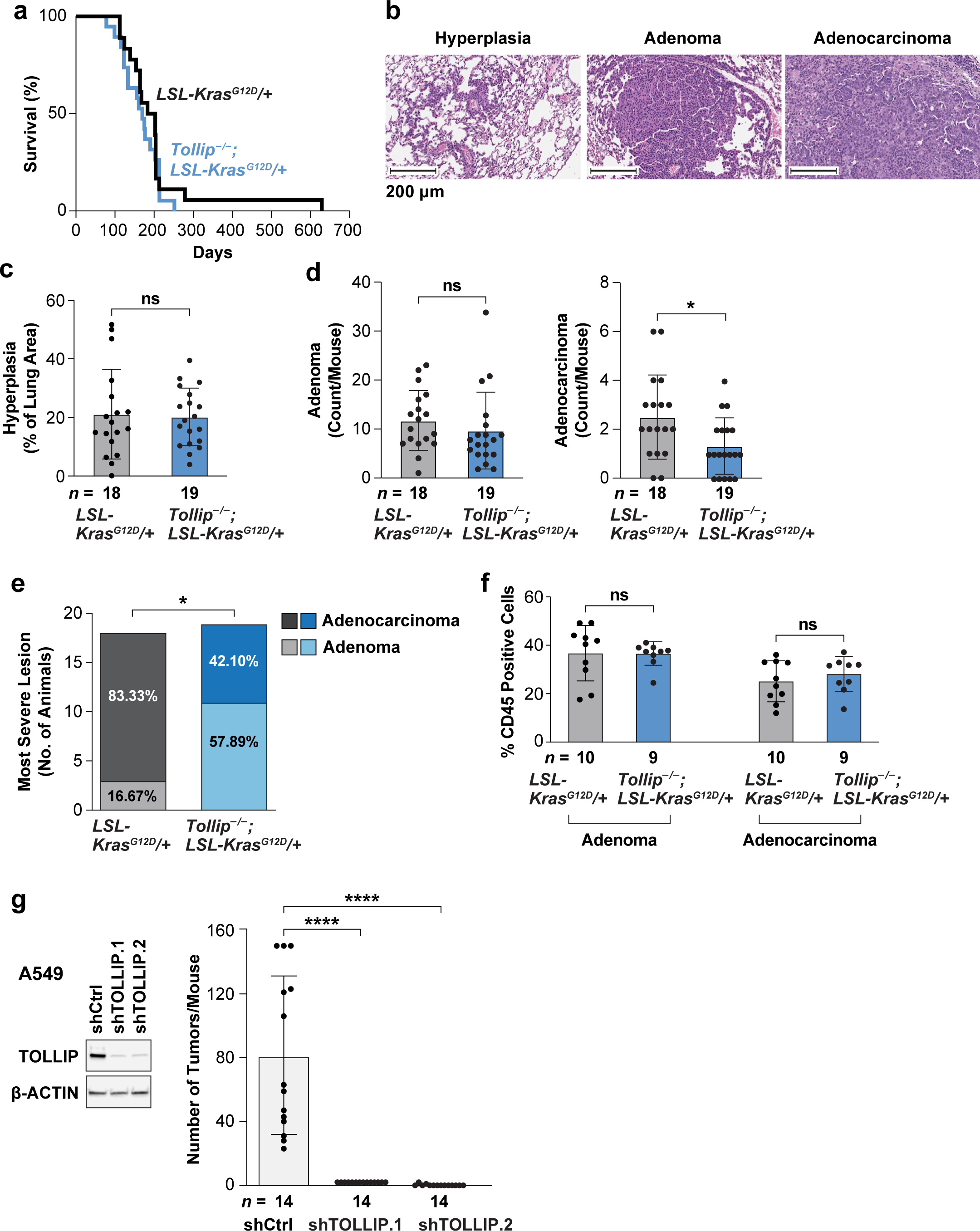
Tollip is required for progression to adenocarcinoma in a *Kras^G12D^*-driven model of lung tumorigenesis. **a** Kaplan-Meier survival curves for *Tollip^+/+^;LSL-Kras^G12D^*and *Tollip^−/−^*;*LSL-Kras^G12D^* mice. Ad.Cre virus was delivered by intratracheal instillation in 10-week-old mice to initiate lung tumorigenesis. Virus administration corresponds to day 1 of the survival study. *Tollip^+/+^*, *n* = 18; *Tollip^−/−^*, *n* = 19. **b** Representative H&E images for lung hyperplasia and adenoma (benign) and ADC (malignant) lesions from *Tollip^+/+^* animals. **c** Hyperplasia burdens in the two strains at clinical sacrifice, measured by HALO image analysis and plotted as % lung area. **d** Adenoma and ADC lesions/mouse, as determined by histopathological examination. **e** Most severe tumor grade observed in each genotype. Scoring is based on histopathological examination of lung lesions. **f** Immune infiltration in lung lesions, as determined by IHC staining for the pan-immune cell marker CD45 in adenomas and ADCs. Data represent % CD45 positive cells in each type of lesion, per animal. **g** Lung nodules in athymic *nu/nu* mice 11 weeks after tail vein injection of 1×10^6^ A549 cells. Cells were infected with a control lentivirus (shCtrl) or two different TOLLIP shRNA vectors prior to injection. TOLLIP knockdown cells produced a maximum of two lung nodules per animal. Left: immunoblots showing TOLLIP knockdown efficiency. The Mantel-Cox log-rank test was used to evaluate Kaplan–Meier survival data. Statistical significance for tumor burdens (**c**, **d**, **f**, **g**) was calculated using two-tailed unpaired Student’s *t*-test. Fisher’s exact test was used to compare the most severe lesion data (**e**). \**P* ≤ 0.05, \*\*\*\**P* ≤ 0.0001.

### Phosphoproteomic analysis identifies targets of perinuclear CK2 and other kinases

The presence of oncogenic kinases such as CK2 on perinuclear endosomes suggests they access a specific set of substrates at this location to generate a pro-oncogenic phosphoproteome. Thus, we sought to identify phospho-sites (p-sites) that require TOLLIP or otherwise correlate with PSC formation. We first characterized TOLLIP-dependent modifications by comparing the phosphoproteomes in control and TOLLIP-depleted A549 cells. Phosphopeptides (p-peptides) were isolated by TiO_2_ enrichment and analyzed by LC-MS/MS. Intensities of individual p-sites were determined by summing the values for all p-peptides that contain a particular p-site and normalizing to total levels of each protein (Fig. 6a). Altogether, 1513 normalized p-sites were detected (Supplementary Table 4).

**Fig. 6.**
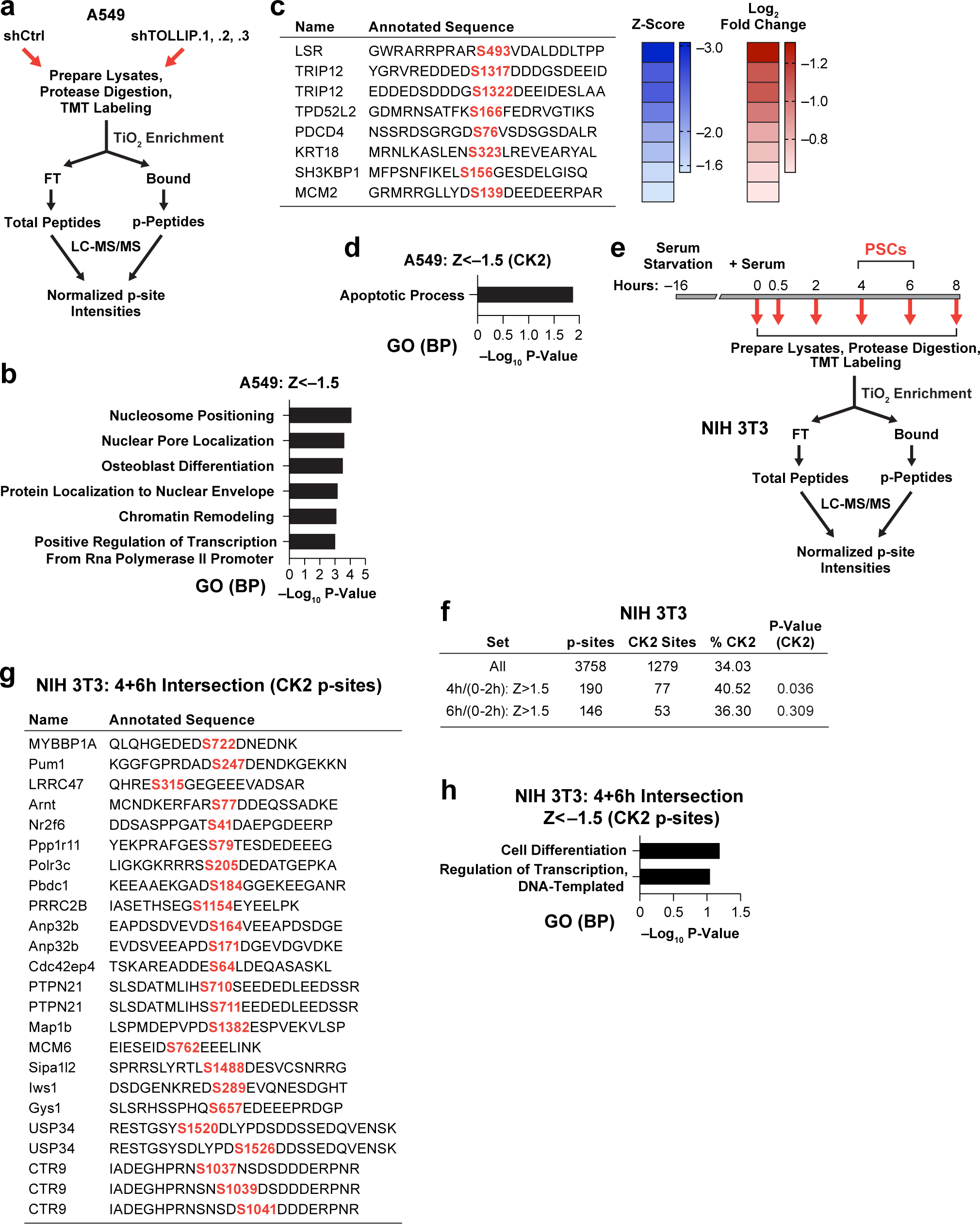
Changes in phosphoproteomes upon TOLLIP depletion in A549 cells or coinciding with PSC formation in GF-stimulated cells. **a** Phosphoproteome analysis pipeline for A549 cells without and with TOLLIP depletion. **b** Ranked gene ontology categories (GO, biological process) for proteins corresponding to p-sites down-regulated in TOLLIP-depleted cells (Z≤-1.5). **c** Sequences of predicted CK2 motifs corresponding to down-regulated p-sites. Right: heat maps showing Z-scores and log_2_ fold changes for each p-site. **d** GO analysis of proteins corresponding to TOLLIP-dependent CK2 p-sites listed in **c**. **e** Phosphoproteome analysis pipeline for serum-stimulated NIH 3T3 cells. KSR1 and CK2α form PSCs at 4 and 6 h, while p-ERK becomes perinuclear at 6 h^17^. **f** Frequency of CK2 motifs in p-sites that increase at 4 and 6 h in NIH 3T3 cells (Z≥1.5) compared to all p-sites. **g** List of common CK2 p-sites up-regulated at both 4 and 6 h. **h** Ranked GO categories (biological process) for common CK2 p-sites shown in **g**. Statistical significance of CK2 site enrichment in NIH 3T3 4 h Z<-1.5 p-sites relative to all p-sites (**f**) was assessed using the one-tailed Binomial test.

Ratios of p-site intensities in TOLLIP-depleted vs. control cells were used to generate Z-scores for each p-site. We applied a Z<-1.5 cutoff, as has been used previously^41^, to define a set of 109 TOLLIP-dependent p-sites (Supplementary Table 4). IPA analysis of these sites revealed one network containing a CK2 node and another with a CDK node (Supplementary Fig. 6a; Supplementary Table 5). GO analysis (Biological Process) showed that the 109 corresponding proteins clustered functionally around chromatin organization and transcription (Fig. 6b; Supplementary Table 6). The Z<-1.5 p-sites were also analyzed for similarity to CK2 phosphorylation motifs and were considered matches if they passed at least one of two rules specifying acidic residues in the surrounding sequence^41,42^ (see Methods). Eight sites scored as putative CK2 motifs (Fig. 6c; Supplementary Table 4). These proteins were functionally enriched for apoptosis (SH3KBP1 and PDCD4) (Fig. 6d; Supplementary Table 6), consistent with increased death of TOLLIP-depleted cells (Fig. 2c).

We also noted a high frequency of CDK motifs in TOLLIP-dependent p-sites. Of the 1513 p-sites in the total A549 data set, 158 (10.44%) were predicted to be CDK sites, while 30 of the 109 Z<-1.5 p-sites (27.52%) scored as potential CDK motifs (Supplementary Fig. 6b,c; Supplementary Table 4). The TOLLIP-dependent Z<-1.5 p-sites were enriched for putative CDK targets when compared to all p-sites (P=5.18E-07). The corresponding proteins are functionally associated with chromatin structure and transcription (GO BP) (Supplementary Fig. 6d; Supplementary Table 6). It remains to be determined whether CDK:cyclin complexes are present on TOLLIP-dependent perinuclear endosomes, or if CDK p-sites are down-regulated through an indirect effect of TOLLIP knockdown.

As an alternative approach to identifying substrates of perinuclear kinases, we used serum-stimulated NIH 3T3 cells to exploit the temporal features of growth factor (GF)-induced PSC formation. In this model, CK2α becomes perinuclear at 4 and 6 h post-stimulation while p-ERK re-localizes at 6 h^17^ (Fig. 6e). Phosphoproteomic analysis was performed on NIH 3T3 lysates harvested at 0, 0.5, 2, 4, 6 and 8 h after GF stimulation and normalized p-site intensities were determined (Supplementary Table 7). To identify p-sites that increase at 4 and/or 6 h, the values at these time points were divided by the combined average of p-site intensities at 0, 0.5 and 2 h. Applying a Z>1.5 threshold revealed 190 p-sites increasing at 4 h and 146 at 6 h (Supplementary Table 7; Supplementary Fig. 6e); these p-sites were significantly enriched for predicted CK2 sites (Fig. 6f). IPA networks based on the 4 h data showed one with a CK2 node and one with a CDK node (Supplementary Fig. 6f; Supplementary Table 8). Filtering for predicted CK2 p-sites revealed 77 sites upregulated at 4 h and 53 at 6 h, of which 27 were common (Fig. 6g; Supplementary Table 7). Considering all p-sites with Z>1.5, the corresponding proteins at both 4 and 6 h were enriched for mRNA processing and RNA splicing (Supplementary Fig. 6g,h; Supplementary Table 9). The 4 and 6 h CK2 p-site-containing proteins also showed associations with RNA processing and rRNA transcription (Supplementary Fig. 6i,j). These results indicate that perinuclear CK2 signaling endosomes appearing during mid-G1 phase in normal cells phosphorylate proteins involved in RNA metabolism, protein synthesis and chromatin structure/transcription.

### TOLLIP-dependent perinuclear CK2 phosphorylates the atypical kinase RIOK1 and controls its pro-oncogenic activity

One of the proteins identified as a likely substrate for perinuclear CK2 is the non-canonical protein kinase, RIOK1. RIOK1 was phosphorylated on a predicted CK2 site, SpS^22^DSE, at 4 h in GF-stimulated NIH 3T3 cells (Supplementary Table 7). RIOK1 plays a key role in ribosome biogenesis and is required for processing of the 18S-E pre-rRNA to the mature 18S form^43^. It also controls assembly/maturation of the 40S ribosomal subunit^44,45,46^ and thus is an important regulator of protein synthesis. *RIOK1* is a synthetic lethal gene in *KRAS* mutant cancers^47^ and is essential for viability of various tumor cell lines^48^ and cancers^49^. Moreover, RIOK1 Ser22 has been implicated as a CK2 site^50^ and is conserved among mammalian RIOK1 homologs (Fig. 7a). Thus, we asked whether pSer22 is a functionally relevant target of perinuclear CK2 in tumor cells.

**Fig. 7.**
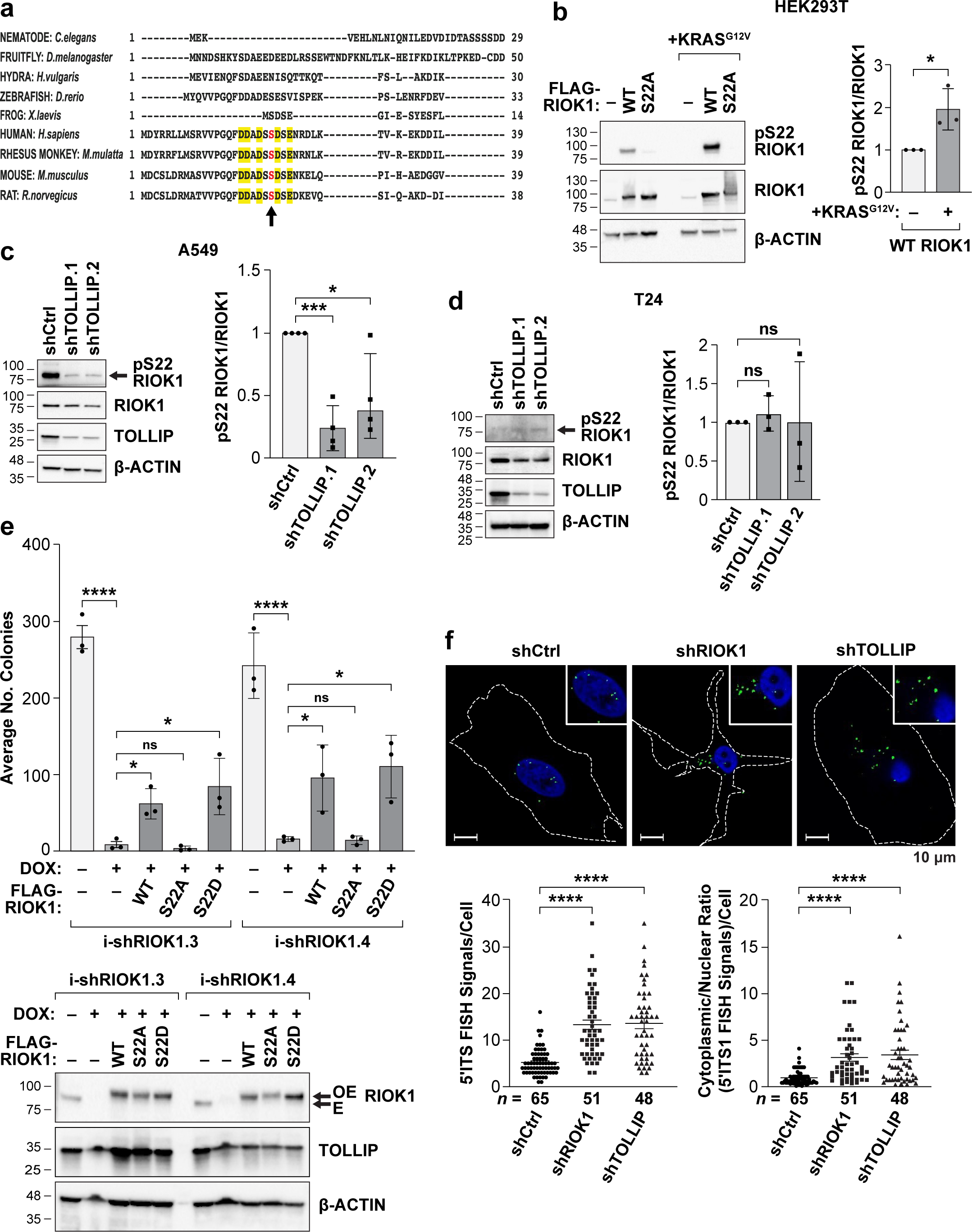
TOLLIP-dependent phosphorylation on RIOK1 Ser22, a CK2 site, stimulates its pro-oncogenic function in tumor cells. **a** RIOK1 Ser22 (arrow) and flanking sequences are conserved in mammals and absent in other species homologs, including other vertebrates. Acidic residues contributing to the CK2 motif are highlighted. **b** RIOK1 Ser22 phosphorylation is augmented by co-expression of KRASG12V. Flagged-tagged RIOK1 WT and S22A constructs were expressed in HEK293T cells, ± KRASG12V. Lysates were analyzed by immunoblotting using a pSer22 RIOK1 antibody. pSer22 RIOK1 levels normalized to total RIOK1 are shown on the right. Data are mean ± s.d.; *n* = 3 independent experiments. **c** TOLLIP depletion decreases RIOK1 Ser22 phosphorylation in A549 cells. RIOK1 pSer22 levels normalized to total RIOK1 are shown on the right. Data are mean ± s.d.; *n* = 3 independent experiments. **d** RIOK1 Ser22 phosphorylation in *HRAS* mutant T24 cells is unaffected by TOLLIP knockdown. pSer22 RIOK1 levels normalized to total RIOK1 are shown on the right. Data are mean ± s.d.; *n* = 3 independent experiments. **E** Ser22 is required for RIOK1 function in A549 cells. RIOK1 was depleted in A549 cells using two different inducible shRIOK1 constructs targeting the 3’UTR. *WT* and S22A and S22D mutant constructs lacking the 3’UTR were expressed in RIOK1-silenced cells to assess their ability to rescue clonogenic growth. Immunoblot shows depletion of endogenous RIOK1 (E) and levels of over-expressed FLAG-RIOK1 proteins (OE). Colony data are mean ± s.d.; *n* = 3 independent experiments. **f** TOLLIP and RIOK1 are required for efficient 18S rRNA processing. A549 cells were depleted for RIOK1 or TOLLIP and analyzed by RNA FISH using a 5’ITS probe to detect the unprocessed 18S precursor. Insets depict enlarged areas encompassing the nucleus. Total number of FISH signals and the cytoplasmic/nuclear ratio per cell were quantified (lower panel); *n*, number of cells analyzed. Two-tailed unpaired Student’s *t*-test was used to analyze pSer22 RIOK1 levels and 5’ITS1 RNA FISH data; \**P* ≤ 0.05, \*\*\**P* ≤ 0.001, \*\*\*\**P* ≤ 0.0001.

To monitor RIOK1 phosphorylation we generated an affinity-purified pSer22 antibody. The antibody recognized over-expressed WT (Ser22) RIOK1 but not S22A, and pSer22 levels increased approximately two-fold upon co-expression with KRAS^G12V^ (Fig. 7b). Endogenous pSer22 was detected in A549 cell lysates and antibody recognition was abolished by incubating lysates with λ-phosphatase (Supplementary Fig. 7a) or treating cells with the CK2 inhibitor, CX-4945 (Supplementary Fig. 7b). Efficient phosphorylation required TOLLIP, as pSer22 signals in A549 cells were reduced by at least 60% following TOLLIP depletion (Fig. 7c). Similar results were observed in PANC1 cells (Supplementary Fig. 7c). Conversely, TOLLIP ablation had no effect on pSer22 levels in *HRAS* mutant T24 cells (Fig. 7d), consistent with the retention of perinuclear CK2 upon TOLLIP knockdown in these cells. These results indicate that Ser22 is an authentic CK2 site whose phosphorylation requires CK2 PSCs.

A549 cells displayed acute dependence on RIOK1 in clonogenic growth assays (Fig. 7e), providing a system for assessing the biological activity of RIOK1 Ser22 mutants. Expression of exogenous WT RIOK1 partially restored colony formation, while S22A did not. However, the phosphomimetic S22D variant rescued RIOK1-depleted cells at least as efficiently as WT RIOK1, indicating that Ser22 phosphorylation stimulates RIOK1 activity to support cell growth/survival.

RIOK1-dependent 18S-E pre-rRNA processing can be monitored by analyzing levels of the 5’ internal transcribed spacer 1 (5’ITS1) in the precursor RNA. When processing is impaired, 18S-E precursors accumulate in the cytoplasm^44^ (Supplementary Fig. 7d). Using RNA FISH, we confirmed that RIOK1 depletion in A549 cells increased cytoplasmic 5’ITS1 signals (Fig. 7f). TOLLIP knockdown similarly augmented cytoplasmic 5’ITS1 levels, comparable to the effect of RIOK1 depletion. These findings support a pathway whereby TOLLIP-dependent, CK2-mediated phosphorylation on RIOK1 Ser22 is necessary to facilitate 18S rRNA processing and 40S ribosome biosynthesis, contributing to the fitness of *K/NRAS* mutant tumor cells.

## Discussion

This study demonstrates that tumor cells require translocation of signaling endosomes to the perinuclear ER network. RAB11A^+^ endosomes containing CK2 and KSR1, but not ERK1/2, rely on TOLLIP for partitioning to the ER. In TOLLIP-depleted tumor cells, CK2, KSR1 and RAB11a revert to a dispersed subcellular distribution resembling that of non-transformed cells, indicating that perinuclear targeting of CK2 and other kinases is a key feature of cancers. TOLLIP was originally identified as a negative regulator of pro-inflammatory signaling^33^ but has not been linked to the RAS pathway in tumor cells. Our demonstration that the ER tethering function of TOLLIP^20^ is essential for oncogenic signaling in neoplastic cells uncovers an important new function for this adapter beyond its currently known roles^39,51,52,53^.

TOLLIP recruits KSR1:CK2 complexes to endosomes by binding to KSR1 through a short, conserved region adjacent to the CUE domain that forms a predicted β-hairpin. While the function of this “linker” region was previously unknown, it was recently shown to mediate the association of TOLLIP and STING (stimulator of interferon genes), stabilizing STING and maintaining cGAS-STING signaling in response to microbial pathogens^54^. It will be of interest to determine if binding of KSR1 and STING to the TOLLIP “linker” region is mutually exclusive and whether TOLLIP’s role in inflammatory signaling is mechanistically distinct from perinuclear targeting of CK2/KSR1 complexes in cancer cells.

The endosome:ER tethering function of TOLLIP is critical for CK2 signaling in a subset of tumor cells, whereas in non-transformed cells PSCs form transiently in response to GF stimulation^17^ and cell proliferation/viability is largely unaffected by TOLLIP depletion. *KRAS*- and *NRAS*-transformed cells, irrespective of the *K/NRAS* activating mutation, are highly dependent on TOLLIP for proliferation and survival. In contrast, tumor cells carrying mutant *HRAS*, *BRAF* and other oncogenes are unaffected by TOLLIP loss. The latter cells nonetheless display perinuclear partitioning of CK2 and KSR1 that is impervious to TOLLIP depletion. These findings point to the existence of alternative pathways for PSC formation involving different endosomal adaptors. Adding further complexity, ERK1/2 kinases are associated with a distinct class of signaling endosomes that lack TOLLIP, with ERK remaining perinuclear in TOLLIP-depleted cells. Moreover, a recent study showed that EGF stimulation causes PI3K to migrate intracellularly along microtubules, leading to AKT activation at perinuclear endosomes rather than at the plasma membrane^55^. Thus, it is possible that active AKT also resides on ER-anchored endosomes. Future work will seek to identify other endosomal oncogenic kinases and adaptors that establish the full spatial signaling landscape in tumor cells, including the adapter that acts redundantly to TOLLIP in non-*K/NRAS* mutant cancers. We observed a pronounced decrease in CDK p-sites upon TOLLIP silencing, raising the possibility that certain CDKs may be associated with TOLLIP^+^ endosomes. However, this result must be interpreted with caution as it could arise from an upstream cell cycle block that secondarily prevents CDK activation. Further studies are needed to determine whether active CDKs are present on perinuclear endosomes.

Our findings implicate PSCs as critical signaling engines in cancers, generating an onco-phosphoproteome required to meet the demands of increased cell proliferation, altered metabolism and pro-apoptotic stresses. p-sites whose phosphorylation corresponds to perinuclear CK2 were found in proteins regulating translation (eIF4B and PDCD4) and ribosome biogenesis (RIOK1 and nucleolin). Hyperactivation of ribosome biogenesis occurs in cancers and can be driven by oncogenes^56^. Accordingly, oncogenic Kras stimulates the RNA-binding activity of nucleolin through phosphorylation by CK2 on four sites^57^, two of which coincide with our GF-stimulated NIH 3T3 data (Supplementary Table 7). These modifications augment the ability of nucleolin to stabilize pre-rRNA transcripts, enhancing ribosome biogenesis. The final step of pre-18S rRNA processing is dependent on RIOK1^43,45,46^ and also requires TOLLIP, and the RIOK1 CK2 site (Ser22) was critical for its ability to support proliferation/survival of A549 cells. Taken together, these observations support a model in which ER-tethered CK2 signaling endosomes specify phosphorylation of pro-oncogenic targets such as RIOK1 and nucleolin in tumor cells (Supplementary Fig. 8). To our knowledge, the present study provides the first definitive evidence that subcellular localization of endosome-associated kinases controls their substrate selectivity.

Tollip facilitates the development of *Kras^G12D^*-driven lung ADCs but is dispensable for benign lesions such as neoplasia and adenomas. Progression to carcinoma in *Kras* lung cancer models has been linked to increased MEK-ERK signaling, with high p-ERK levels often associated with amplification of the mutant *Kras* locus^58^. Interestingly, *Kras*-induced adenomas display perinuclear CK2, KSR1^17^ and Tollip (Supplementary Fig. 1g) even though these benign lesions develop efficiently in the absence of Tollip. Evidently there is a transition to Tollip dependence that accompanies progression to lung ADC, possibly involving enhanced RAS pathway signaling. Human *KRAS* mutant cell lines are nearly always derived from late-stage malignant carcinomas, which likely explains their TOLLIP addiction. As TOLLIP is an Achilles heel of *KRAS* and *NRAS* tumor cells, it could potentially be exploited as a therapeutic target in these cancers.

## Acknowledgements

We thank Dr. A. Ciarrocchi for providing *Tollip* constructs, T.N. O’Sullivan for generating KPC pancreatic tumor samples, and A. Kane (Scientific Publications, Graphics & Media, Leidos Biomedical Research, Inc., Frederick National Laboratory for Cancer Research) for assistance with figure preparation. This research was supported in part by the Intramural Research Program of the NIH, National Cancer Institute, Center for Cancer Research, and in part with Federal funds from the National Cancer Institute, National Institutes of Health, under Contract Nos. HHSN261200800001E and 75N91019D00024. Analysis and management of images and associated metadata was supported in part by the NCI HALO Image Analysis Resource. The content of this publication does not necessarily reflect the views or policies of the Department of Health and Human Services, nor does mention of trade names, commercial products, or organizations imply endorsement by the U.S. Government.

## Contributions

S.B. and P.F.J. conceived the study and designed experiments. S.B. performed the experiments. B.T.L. and M.G. provided bioinformatic analysis of phosphoproteomic and RNA-seq data, respectively. K.S. and N.M. conducted animal experiments and genotyping. S.K. provided mouse pancreatic tumor samples. B.K. and L.B. performed histopathological analysis of lung tumor samples. S.L. devised the method to calculate RNPI and implemented it in a Fiji macro script. S.D. and T.A. designed and performed mass spectrometry experiments and data analysis. D.E. provided key plasmid constructs. S.B. and P.F.J. wrote the manuscript with input from all authors.

## Methods

### Animal experiments

Mice were maintained in accordance with NIH animal guidelines following protocols approved by the NCI-Frederick Animal Care and Use Committee. Mouse embryonic fibroblasts (MEFs) were isolated from *WT*, *Tollip^−/−^*^53^*, p19^Arf−/−^* and *p19^Arf−/−^;Tollip^−/−^* E13.5 mouse embryos (C57Bl/6Ncr strain background), and cells were maintained at low passage. *Kras^LA^*^2^/+^59^, *LSL-Kras^G12D^/+*^60^ and *Tollip^−/−^;LSL-Kras^G12D^/+* mice (C57Bl/6Ncr background) were used for tumor studies. To induce lung tumors in *LSL-Kras^G12D^* strains, Ad.Cre virus (Viral Vector Core Facility, University of Iowa College of Medicine) was administered by intratracheal instillation of 1.5 or 3.0×10^7^ pfu/animal. Animals that became moribund (thin, hunched, labored breathing, rough coat, or lethargy) were sacrificed, lungs were removed and fixed in 10% Neutral Buffered Formalin (NBF) and embedded in paraffin blocks. Lungs from normal mice were used as controls. NBF fixed paraffin-embedded pancreatic tumor tissue sections were obtained from KPC mice (*Pdx-1*-*Cre (B6.Tg(Pdx1-cre)6Tuv/Nci*); *LSL-Kras^G12D^* (*B6.Krastm4Tyj/Nci*); *LSL p53^R^*^172^*^H^ (129S4-Trp53tm2Tyj/Nci)* on a C57BL/6 background)^61^. Pancreas samples from *WT* mice were used as controls. Primary tumor cell line KPC98027 was derived from a KPC pancreatic ductal ADC. 1×10^5^ KPC cells were injected into medial pancreata in syngeneic C57BL/6 recipient animals. Pancreas from un-injected normal mice were used as controls. Tumors were collected at 5-6 weeks post-injection and fixed in NBF as described above. For xenograft experiments, athymic nude (*nu/nu*) mice received 1×10^6^ A549 cells (control or TOLLIP knockdown) by tail vein injection. All mice were sacrificed when a single mouse showed clinical signs and became moribund. Lungs were removed and lesions counted. Lungs were gently inflated in NBF, fixed with NBF for 5 days and embedded in paraffin blocks or stored in 70% ETOH.

### H&E staining, tumor grading and HALO AI-based image analysis of lung tumor slides

Mouse lungs were formalin-fixed, embedded in paraffin and sectioned at 5µm thickness. Routine hematoxylin and eosin-staining was performed using the Sakura® Tissue-Tek® Prisma™ automated stainer (Sakura Finetek USA, Inc., Torrance, California). The slides were dewaxed using xylene and then hydrated using a series of graded ethyl alcohols. Commercial hematoxylin, clarifier, bluing reagent and eosin-Y were used to stain. A regressive staining method was used, which intentionally overstains tissues and then uses a differentiation step (clarifier/bluing reagents) to remove excess stain. After staining was completed, the slides were cover slipped using the Sakura® Tissue-Tek™Glass® automatic cover slipper (Sakura Finetek USA, Inc., Torrance, California). Whole slide images were obtained at high resolution, 20x magnification using an AT2 scanner (Aperio, Leica Biosystems, Buffalo Grove, IL, USA).

Two veterinary pathologists (B.K. and L.B.) identified and segmented the regions of interest used to train an AI deep learning tissue-classifying algorithm (Densenet AI V2 plugin in Halo (v.3.3.2541.423, Indica Labs, Corrales, NM, USA). Annotations were manually drawn around four tissue classes: normal (including lung parenchyma, airways and vessels), hyperplasia (hypercellular foci of round to cuboidal hypertrophic pneumocytes arranged in a single layer), adenoma (expansile mass consisting of proliferative epithelium arranged in papillary to solid formations with no features of atypia and rare to absent mitotic figures), and adenocarcinoma (proliferative cells demonstrating some combination of increased cellular atypia, anisocytosis, anisokaryosis, cytoplasmic basophilia, increased mitotic figures and haphazard or invasive patterns).

During training, the algorithm performance was iteratively evaluated by visual inspection and misclassification of tissues was addressed and corrected by additional annotations in the misclassified regions and further training until satisfactory performance was obtained. The percentages of lungs comprised of hyperplasia, adenoma and adenocarcinoma were calculated by dividing the total area of each class by the total tissue area on the slide and multiplying by 100.

### Immunohistochemistry

Immunohistochemical staining for CD45 was performed on 5-micron FFPE sections using a manual benchtop method. Antigen retrieval was performed with citrate buffer for 30 seconds at 121°C using a Decloaking Chamber (BioCare Medical). Nonspecific binding was blocked with an incubation of 2% normal rabbit serum for 20 min. This was followed by an overnight incubation at 4°C with the CD45 primary antibody (BD Biosciences #550539) at a 1:100 dilution. Antibody detection was accomplished by incubation with Biotinylated Rabbit anti-Rat (Vector labs# BA-4001) secondary antibody and ABC Elite reagent (Vector Labs # PK-6100), followed by incubation in DAB chromogen (Sigma #D5905). Slides were digitally scanned at 20x using a Leica AT2 whole slide scanner. Normal mouse spleen was used a positive control.

Regions having adenoma and adenocarcinoma lesions were graded as mentioned above using Halo AI on the H&E slides, and CD45 staining in the corresponding regions was analyzed. Hematoxylin nuclear stain was used as a marker to count the total number of cells within the graded adenoma or adenocarcinoma lesions. The percentage of CD45 stained cells was plotted.

### Cells and cell culture

HEK-293T cells (American Type Culture Collection (ATCC), Rockville, MD) and MEFs were cultured in DMEM+Glutamax (supplemented with 10% Fetal Bovine Serum (FBS). HBEC cells (ATCC) were cultured in Airway Epithelial Cell Basal Medium (ATCC® PCS-300-030) supplemented with Bronchial Epithelial Cell Growth Kit (ATCC PCS-300-040). NIH3T3 cells were grown in DMEM+Glutamax with 10% calf serum (Colorado Serum Company). A549 (ATCC) cells were grown in Ham’s F-12K (Kaighn’s) medium with 10% FBS. PANC-1, A375, MDA-MB-231, HepG2 and MIA PaCa-2 cells (ATCC) were grown in DMEM+Glutamax with 10% FBS. PC-3, OVCAR-8 and NCI-H1299 cells (ATCC) were grown in RPMI+Glutamax with 10%FBS. KPC98027 cells were cultured in DMEM/F12+Glutamax with 10% FBS. *HRAS* mutant cancer cell lines T24 and RL95-2 were kindly provided by Dr. Deborah Morrison (NCI-Frederick) and were grown in McCoy’s 5A and DMEM/F12+Glutamax, respectively. HT-29 cells were cultured in McCoy’s 5A medium with 10% FBS. All media and FBS were obtained from Thermo Fisher Scientific. To prevent Mycoplasma contamination, cell cultures were additionally supplemented with Plasmocin (InvivoGen). Cell stocks were generally passaged fewer than 5 times, and freshly thawed cells were maintained in culture for no more than 2 weeks before conducting experiments.

### Antibodies

Rabbit polyclonal antibodies to TOLLIP (11315-1-AP) and RAB11A (20229-1-AP) were obtained from Proteintech Group. Mouse monoclonal antibody to Tollip (Clone: MAB4678) was obtained from R&D Systems. Rabbit Anti-CSNK2A1 (CK2α) antibody (ab10466) was from Abcam. Rabbit antibodies to phospho-ERK1/2 (#9101) and HA-Tag (#3724) were from Cell Signaling Technologies. Mouse antibodies to β-Actin (sc-47778) and p-ERK1/2 (sc-136521) were obtained from Santa Cruz Biotechnology. Rabbit antibodies to HRAS (GTX-116041) and KSR1 (GTX56241) were from GeneTex. Affinity purified mouse antibody against Pyo epitope tag (Glu-Glu) was from BioLegend (#901802). Mouse monoclonal antibody against KRAS (H00003845-M01) was obtained from Abnova. Anti-mouse (W4028) and anti-rabbit (W4018) HRP conjugated secondary antibodies were from Promega. Anti-mouse Alexa® Fluor 488 (#A32790) and anti-rabbit Alexa® Fluor 594 (A-21207) conjugated secondary antibodies were from Thermo Fisher Scientific. A custom phospho-Ser22 RIOK1 polyclonal antibody (Biosynth) was raised in rabbits using the immunogenic peptide H-Cys-Asp-Asp-Ala-Asp-Ser-**p**Ser-Asp-Ser-Glu-Asn-NH2. Peptides were synthesized and purified using high-performance liquid chromatography (HPLC) to >90% purity as determined by HPLC and/or LC/MS. The immunogen was conjugated to activated keyhole limpet hemocyanin (KLH). Two Specific Pathogen Free New Zealand white rabbits (12-16 weeks old) were immunized subcutaneously with the phospho-peptide conjugate. All animal protocols were approved by the Institutional Animal Care and Use Committee. To remove any latent “non-phospho” reactive antibodies, the anti-sera were passed twice through SulfoLink resin (ThermoFisher scientific # 20402) columns coupled to the non-phosphorylated peptide. The final flowthrough was passed through SulfoLink resin coupled to the phosphorylated peptide to obtain affinity-purified phospho-RIOK1 Ser-22 antibodies.

### Reagents

1 mg/ml stock solution of Doxycycline hyclate (Sigma Aldrich, # D9891-1g) were prepared in distilled water and filter sterilized. The final concentration of Doxycycline (Dox) used was 1µg/ml. CK2 inhibitor, CX-4945 (Silmitaserib) was purchased from Activate Scientific (#AS10115-G1) and λ-phosphatase (λ-PPTase) was obtained from Merck Millipore Sigma (#P9614-20KU). Thapsigargin (T9033, Millipore Sigma) was used at 400nM concentration.

### Plasmids and lentiviral vectors

*Ksr1* expression plasmids containing an N-terminal Pyo tag (EYMPME) were generated by PCR amplification using pcDNA3 mouse *Ksr1*^62^ as a template and inserting fragments into pcDNA3.1. C-terminal deletion mutants 1-81 (CA1), 1-286 (CA1-2) were cloned using EcoRI and BamH1 sites, while 1-377 (CA1-3), and 1-479 (CA1-4) were cloned using EcoR1 and Kpn1 restriction sites. N-terminal deletion mutant 531-873 (CA5) was inserted using EcoR1 and BamH1 sites. Expression plasmids for HA-tagged WT rat *Tollip* and deletion mutants (Δ229-274, Δ180-274, Δ1-54 and Δ1-94) in pRK7^32^ were a kind gift from Dr. A. Ciarrocchi (Laboratory of Translational Research, Azienda Unità Sanitaria Locale-IRCCS di Reggio Emilia, Italy). HA-*Tollip* deletion mutant Δ203-274 was made by PCR amplification and cloning into the pRK7 vector using BamH1 and EcoR1 restriction sites. Δ180-202 was made using three-way ligation of cDNA fragments encoding *Tollip* residues 1-179 and 203-274, and BamH1/EcoR1 digested pRK7 vector; Nhe1 sites were used to join the two *Tollip* cDNA fragments. pLenti CMV GFP Hygro (656-4) was a gift from Eric Campeau and Paul Kaufman (Addgene plasmid #17446; http://n2t.net/addgene:17446; RRID:Addgene_17446)^63^. Rat WT *Tollip* and deletion mutants Δ229-274, Δ180-274, Δ1-54, Δ180-202 were cloned into pLenti CMV GFP Hygro using PCR amplification from the pRK7 based Rat *Tollip* plasmids mentioned above, using primers containing BsrG1 and Sal1 sites. Δ55-179 was made using three-way ligation of cDNA fragments encoding *Tollip* residues 1-54 and 180-274, and BsrG1/Sal1 digested pLenti CMV GFP Hygro vector; Nhe1 sites were used to join the two *Tollip* cDNA fragments. Human *TOLLIP* was amplified from cDNA isolated from A549 cells using primers containing BsrG1 and Sal1 sites and the product inserted into pLenti CMV GFP hygro vector. A BsrG1 site within the *TOLLIP* cDNA was eliminated by mutation while maintaining the amino acid sequence, using the QuikChange Lightning Site-Directed Mutagenesis Kit (Agilent, #210518) per the manufacturer’s instructions. Lentiviral Rat *Tollip* constructs containing C-terminal fusion to EGFP were generated using three-way Gateway cloning. The minimal CMV promoter-containing pDONR258 and C-terminal m-EGFP tag-containing pDONR257 were obtained from the NCI RAS Initiative (Leidos Biomedical Research, Inc., Frederick, MD, USA). Coding sequences for Rat *Tollip* WT and deletion mutants were PCR amplified from the corresponding pLenti CMV GFP hygro vectors (described above) and cloned into pDONR255. The CMV promoter, either Rat *Tollip* WT, Δ229–274, Δ180-274, Δ1-54, Δ180-202 or Δ55-179 coding regions, and m-EGFP were then inserted into the lentiviral pDest659 vector (carrying Blasticidin resistance and attR4/attR3 recombination sites). The resulting *Tollip* WT or deletion mutant vectors were used for viral production and transduction. Lentiviral expression vectors for GFP- or mCherry-tagged human *KSR1*, mCherry-tagged mouse *Ck2α*, and FLAG-tagged human *RIOK1* were obtained from Genecopoeia (Rockville, MD). Lentiviral *RIOK1* constructs were generated using four-way Gateway cloning. The minimal CMV promoter-containing pDONR259 vector and the 3X-FLAG tag containing pDONR235 vector were obtained from the NCI RAS Initiative (Leidos Biomedical Research, Inc., Frederick, MD, USA). The human WT *RIOK1* coding region was PCR amplified from cDNA isolated from A549 cells; oligos containing the RIOK1 polyadenylation signal (PAS) were annealed. *RIOK1* coding region and *RIOK1* PAS fragments were then cloned into pDONR255 and pDONR257 vectors, respectively, using BP recombination. The *RIOK1* Ser22 to Ala (A) or Ser22 to Aspartate (D) mutants were generated using QuikChange Lightning Site-Directed Mutagenesis Kit (Agilent, Catalog #210518). Primers were designed using the QuikChange Primer Design website. Using four-way LR recombination, the CMV promoter, 3X-FLAG, either WT, S22A or S22D *RIOK1* coding regions, and the *RIOK1* PAS fragment were cloned into the lentiviral pDest659 vector. The resulting *RIOK1* WT, S22A or S22D vectors were used for either transfection or viral production and transduction. Lentiviral vectors for human *HRAS^G12V^* and *KRAS^G12V^* were obtained from the NCI RAS Initiative (Leidos Biomedical Research, Inc., Frederick, MD, USA). Lentiviral packaging/envelope plasmids pMD2.G (#12259), pMDLg/pRRE (#12251), and pRSV/Rev (#12253) were obtained from AddGene. pLV-ER GFP encoding GFP-tagged *SEC61β* was a gift from Pantelis Tsoulfas (Addgene plasmid #80069; http://n2t.net/addgene:80069; RRID:Addgene_80069). mEmerald-RAB11A-7 was a gift from Michael Davidson (Addgene plasmid #54245; http://n2t.net/addgene:54245; RRID:Addgene_54245). pTag-RFP-C-h-Rab11a-c-Myc was a gift from James Johnson (Addgene plasmid #79806; http://n2t.net/addgene:79806; RRID: Addgene_79806)^64^. The lentiviral pUltra-hot vector was a gift from Malcolm Moore (Addgene plasmid #24130; http://n2t.net/addgene:24130; RRID: Addgene_24130). mEmerald tagged human *RAB11A* and RFP tagged human *RAB11A* were transferred into the Ultra-hot vector using Age1 and BamH1 cloning sites. Similarly, mCherry tagged human *TOLLIP* was cloned into the Ultra-hot vector. Mission shRNA lentiviral constructs for human *TOLLIP* (TRCN0000314717, TRCN0000356024 and TRCN0000314642), mouse *Tollip* (TRCN0000066136 and TRCN0000066137), human *RAB11A* (TRCN0000073022 and TRCN0000291939) and human *KSR1* (TRCN0000338414 TRCN0000338476) were obtained from Millipore Sigma. SMARTvector Inducible Human RIOK1 hEF1a-TurboGFP shRNAs [#V3SH11252-224914513 (i-shRIOK1.3) and #V3SH11252-229953481(i-shRIOK1.4)], were obtained from Dharmacon (Horizon Discovery). All oligonucleotide primers used in cloning procedures are shown in Supplementary Table 10.

### Transient transfection

Transfections were carried out using FuGENE 6 transfection reagent (Promega #E2691) per the manufacturer’s instructions. Briefly, HEK293T cells in 100mm dishes were transfected with varying amounts of Pyo-tagged *Ksr1* or deletion mutants (standardized to obtain equivalent protein expression), HA-tagged *Tollip* or *Tollip* deletion mutants (standardized to obtain equivalent protein expression), or 2µg of FLAG-tagged human *RIOK1* or *RIOK1* S22A with or without 0.1µg pcDNA3-*KRAS^G12V^*. Empty pcDNA3 was added when necessary to equalize total vector DNA. 2µl FuGENE 6 transfection reagent per µg of DNA was used for each transfection. Transfection mix was made in antibiotic free Opti-MEM medium (Thermo Fisher Scientific, Catalog #11058021). After transfection, cells were cultured in complete media (DMEM with 10% FBS) for 24 h. Media was changed, and cells were incubated for another 24 h before harvesting. For experiments involving KRAS^G12V^, media was changed 24 h after transfection and cells were incubated for 12 h in complete media followed by overnight incubation in low serum conditions (DMEM with 0.1% FBS) prior to harvesting.

### Lentiviral transduction

Lentiviral plasmids were transfected into 293T cells using a standard CaPO_4_ precipitation method with 20 μg lentiviral vector, 6 μg pMD2.G, 10 μg pMDLg/pRRE and 5 μg pRSV/Rev. After 24 h, cells were washed with phosphate-buffered saline (PBS) and supplemented with fresh media. 48-72 h after transfection, viral supernatants were collected, pooled, filtered through 0.45 mm filters, supplemented with 8 µg/ml polybrene and used to infect target cells. 24 h later, media was removed, cells washed with PBS and supplemented with fresh media and the appropriate antibiotic. Multiple genes were introduced by sequential infection and drug selection.

### Growth curves

Cells were seeded at 2.5×10^4^ cells/well in 6-well dishes. At the appropriate times, cells were washed with PBS, fixed in 10% formalin, rinsed with water, stained with 0.1% crystal violet (Millipore Sigma) for 30 min, rinsed extensively, and dried. Dye was extracted with 10% acetic acid and absorbance measured at 590 nm. All values were normalized to day 0 (24 h after plating). Growth curves were plotted using GraphPad Prism 10.2.0.

### Clonogenic growth assays

4000 cells (A549, PANC1, MDA-MB-231, MIA PaCa-2, H1299, HepG2, RL95-2, A375, HT29, OVCAR-8 and PC3) or 1000 cells (T24) or their respective derivatives were plated on 100 mm culture dishes. After 10-14 days, colonies were fixed with 4% formaldehyde in PBS for 10 min, stained with 0.4% crystal violet for 30 min, rinsed thoroughly with water, dried and scanned. The number of colonies were counted using FIJI/ImageJ software v1.53f51.

### Senescence-associated β-Galactosidase (SA-βGal) assays

A549 cells or their derivatives were plated at 2.5×10^4^ cells per well in 6-well plates. After 2-3 days, cells were stained using a senescence detection kit (Millipore Sigma QIA117) per the manufacturer’s instructions.

### Immunoprecipitation (IP)

1×10^6^ HEK293T cells were seeded in 10 cm dishes and transfected with DNA combinations using FuGENE6 (Promega) as described above. 48 h after transfection, cells were washed twice with PBS and processed for cell lysis. For IP of endogenous TOLLIP and exogenously expressed pyo-KSR1, non-denaturing Lysis Buffer (137 mM NaCl, 20 mM Tris pH 8.0, 10% glycerol,1% Triton X-100) was used. Lysis buffer supplemented with freshly prepared protease inhibitors (Protease Inhibitor Cocktail Set I #539131 and Protease Inhibitor Cocktail Set III #539134, Millipore Sigma-Aldrich) and phosphatase inhibitors (Phosphatase Inhibitor Cocktail Set I #524624 and Phosphatase Inhibitor Cocktail Set II #524625; Millipore Sigma-Aldrich) was added directly to the plated cells for 15 min on ice. Lysates were centrifuged at 13,000 rpm for 5 min, supernatants collected, and protein estimated using Bradford reagent (Bio-Rad Laboratories, #50000006). 2 mg supernatant was used for immunoprecipitation, with 10% reserved for input samples. For IP of endogenous TOLLIP, cell lysates were pre-cleared by adding 30 μl of a 50% slurry of Protein G Sepharose (GE Healthcare, #17-0618-01) for 1 h. For pre-clearing for pyo-KSR1 IP, 10µl of Protein G magnetic beads (NEB, S1430S) was used. Precleared supernatant was collected either by centrifugation at 1000 rpm or using a magnetic rack (NEB). Anti-TOLLIP or anti-Glu Glu (Pyo) antibody (1:1000) was added to the pre-cleared supernatants and incubated for 1 h at 4°C on a rotating rack. 30μl of 50% slurry of Protein G Sepharose or 10µl Protein G magnetic beads were added and incubated at 4°C on a rotating rack for 1 h. Beads were collected and washed 4 times in NET-N Wash Buffer (100mM NaCl, 20 mM Tris pH 8.0, 1 mM EDTA and 0.5% NP40). 2xSDS sample buffer (Bio-Rad Laboratories) was added to the beads and samples boiled for 7 min at 100°C. Proteins were analyzed by immunoblotting as described below.

### Immunoblotting

Cells were harvested after washing twice with cold PBS and lysed with IP lysis buffer or RIPA lysis buffer supplemented with protease and phosphatase inhibitors (see above for details). Cells were lysed on ice for 15 min. Protein samples (30-50 µg) or IP samples were resolved by electrophoresis on 10% or 12% Mini or Midi-TGX Precast Gels (Bio-Rad Laboratories, Cat. nos: 4561034, 4561044, 5671044 and 5671034) and transferred to PVDF membranes (Bio-Rad Laboratories, Cat. no. 1704273). The blots were probed with the appropriate primary antibodies followed by goat anti-rabbit IgG or goat anti-mouse IgG conjugated to horseradish peroxidase and visualized using enhanced chemiluminescence substrate (SuperSignal™ West Dura Extended Duration Substrate #34076, Thermo Fisher Scientific).

### Live cell imaging

1×10^4^ cells were plated in glass-bottomed chambers (Sarstedt Cat. no. 94.6190.402) or μ-Slides VI0.4 (Ibidi) and cultured for 24 h. Cells were then infected with supernatants of the appropriate lentiviral vectors. After 18-24 h, media was replaced, and cells were grown in DMEM+10%FBS for 24h. Media was then replaced with DMEM +0.1% FBS and cells cultured overnight. Cells were imaged using a Zeiss LSM-780 confocal microscope equipped with an incubation chamber maintained at 37°C in 5% CO_2_.

### Immunofluorescence (IF)

1×10^4^ cells were plated in glass-bottomed chambers (Sarstedt, Cat. no. 94.6190.402) or μ-Slides VI0.4 (Ibidi) and cultured for 24 h. Cells were serum starved overnight and fixed with ice cold 100% methanol for 20 min. The fixed cells were incubated for 1 h in blocking buffer [5% Normal Goat Serum (Cell Signaling) in PBS with 0.3% TritonX100 (Sigma)]. Thereafter, the cells were incubated with the appropriate primary antibodies (1:1000 dilution) overnight at 4°C in blocking buffer. Following 4 washes with IF wash buffer (PBS containing 1% BSA and 0.1% TritonX100), Alexa 488 or Alexa 594 conjugated secondary antibodies (1:1000 dilution) in IF wash buffer were added for 1 h at RT. The cells were then washed 4 times and stained with 0.1 μg/mL DAPI (Thermo Fisher Scientific) in PBS for 12 min at room temperature followed by 3 washes with PBS. Fluorescence images were acquired using a Zeiss LSM-780 confocal microscope. For IF of tumor samples, dissected tumor-bearing lungs or pancreata embedded in paraffin blocks (described above); were dissected and 5µm sections were prepared and slides processed for immunofluorescence. Briefly, slides were deparaffinized using xylene, rehydrated by treating with sequentially decreasing concentrations of ethanol, and antigen retrieval performed using sodium citrate buffer (10mM sodium citrate, 0.05% Tween 20 and 1M HCl). Thereafter, the slides were washed with PBST, fixed in ice-cold methanol for 10 min at -20°C, permeabilized with 1% TritonX100 for 15 min at RT and 1% SDS for 10 min at RT; all steps were interspersed with 3 PBS washes. Slides were blocked using 5% goat serum with 0.3% TritonX-100 in PBS, followed by overnight incubation with rabbit primary antibodies targeting Tollip, KSR1, CK2a or p-ERK at 1:500 dilution in blocking buffer, followed by washing with IF wash buffer (0.1% Triton X-100/1% BSA in PBS) and incubating with Alexa 594 conjugated secondary anti-rabbit antibody (1:500 dilution in IF wash buffer) for 1 h at RT. DAPI staining was performed as described above and the slides were mounted on cover slips using Vecta-shield mounting media (VECTASHIELD® Antifade Mounting Medium, H-1000, VectorLabs). Fluorescence images were acquired using a Zeiss LSM-780 confocal microscope.

### Fluorescence-based image analysis

Relative nuclear proximity index (RNPI), which measures the proximity of cytoplasmic fluorescence signals to the nuclear envelope, was used to analyze manually selected lateral (XY) confocal slices using FIJI/ImageJ v1.53f51 image analysis software^65^. Using the Differential Interference Contrast (DIC) image and the DAPI stained nucleus, the cell border was traced manually and the nuclear border was drawn around the cell nucleus. Then a Fiji macro (available with instructions from the authors upon request) performed the proximity analysis. The macro first required the user to set a threshold intensity to discriminate punctate fluorescence signals from background intensity. Then the macro automatically made four calculations: (1) The area of the cytoplasm, (2) The total intensity of the fluorescence signals above the manually set threshold intensity (the macro did not subtract the background intensity from the fluorescence signals.), (3) The closest distance to the nuclear border determined for each pixel in the cytoplasm (excluding the nucleus) using the Euclidean distance transform, and then integration of these distance values over the cytoplasm, and (4) The product of the fluorescence intensity above background calculated for each pixel in the cytoplasm [from (2)] and the distance value [from (3)], and then integration of these products over the cytoplasm. The manually set threshold intensity was kept constant in an individual experimental set. The four values from these calculations were used to calculate the average distance of the fluorescence signals from the edge of the nucleus [the value from (4) divided by the value from (2)], and to calculate the average distance a uniform signal in the cytoplasm would have from the edge of the nucleus, which is the value from (3) divided by the value from (1). The ratio of the average distance of the fluorescence signals from the edge of the nucleus to the average distance a uniform signal in the cytoplasm would have from the edge of the nucleus was calculated for each cell. If the ratio is significantly above 1 then the fluorescence signals are further away from the nucleus than a uniformly distributed signal. If the ratio is significantly below 1 then the fluorescence signals are closer to the nucleus than a uniformly distributed signal. RNPI values were plotted using Graphpad Prism 10.2.0.

Pearson’s correlation coefficient and Manders overlap coefficient (Zen Blue software-based Image analysis tool) were used to determine colocalization between proteins (Pearson’s coefficient value quantifies the degree to which fluorescence pixels from two channels follow a simple linear relationship of intensity, while Manders coefficient quantifies the degree of overlap between two fluorescent pixels). The resultant values were plotted using GraphPad Prism 10.2.0.

### Proximity ligation assay (PLA)

1×10^4^ A549 cells were grown on glass bottom μ-Slides VI0.4 (Ibidi). PLA was performed using the Duolink *in situ* PLA kit according to the manufacturer’s instructions (Millipore Sigma, DUO92002, DUO92004 and DUO92014). In brief, cells were fixed with pre-chilled MeOH for 15 min at -20°C and permeabilized with 0.05% Saponin in PBS. After blocking, the cells were incubated overnight at 4°C with the appropriate combinations of antibodies. PLA Plus and Minus probes for mouse and rabbit antibodies (used at 1:5 dilution in Duolink® Antibody Diluent) were added and incubated for 1 h at 37°C in a pre-heated humidified chamber. The proximity ligation reaction and the polymerase-based DNA amplification reaction were performed for 30 min and 1 h 40 min, respectively, at 37°C. Samples were counterstained with DAPI for 12 min and images were acquired using a Zeiss LSM-780 confocal microscope. PLA spots were quantified using FIJI/ImageJ software v1.53f51.

### RNA FISH

1×10^4^ A549 cells were plated on μ-Slides VI0.4 (Ibidi). 48 h after seeding, cells were washed in Cytoskeleton buffer (CB; 10 mmol/L MES pH 6.1, 150 mmol/L NaCl, 5 mmol/L MgCl_2_, 5 mmol/L EGTA, 5 mmol/L glucose) and permeabilized/fixed in ice-cold pre-fixative mix (4% paraformaldehyde, 0.01% glutaraldehyde, 0.05% saponin, in CB) for 20 min at 4°C. Cells were then incubated with ice-cold fixative mix (4% paraformaldehyde, 0.01% glutaraldehyde, in CB) for 100 min at 4°C. Fixed cells were washed with CB twice and quenched by addition of 50 mmol/L NH_4_Cl in CB for 5 min at RT. Protease digestion (1:2000 dilution) was performed for 10 min at RT using Protease-QS solution in PBS followed by 2 washes with 1X PBS. For RNA FISH, QuantiGene ViewRNA ISH Cell Assay (Thermo Fisher Scientific) was used according to the manufacturer’s protocol. A custom-made probe for human pre-18S rRNA 5’ITS1 (internal transcribed spacer 1) (5′-CCTCGCCCTCCGGGCTCCGTTAATGATC-3′) was used^44^. Cells were washed four times with PBS-S (0.05% Saponin, in PBS) for 5 min at 20°C, counterstained with DAPI for 10 min at RT, and then washed twice with 1X PBS. Images were acquired using a Zeiss LSM-780 confocal microscope. Image analysis was performed using FIJI/ImageJ software v1.53f51. Briefly, cell border and nuclear border were drawn and marked as region of interest (ROI) and the number of particles (mRNA foci) within the ROI were calculated. The number of cytoplasmic signals was calculated as the number of 5’ITS1 FISH signals within the cell border subtracted by the number of signals within the nuclear boundary. The total number of 5’ITS1 FISH signals per cell and the ratio of cytoplasmic to nuclear FISH signals per cell were plotted using GraphPad Prism 10.2.0.

### Cell processing for phospho-proteomic analysis

2×10^6^ NIH 3T3 cells were seeded in 150mm plates and cultured for 24 h in DMEM media containing 10% calf serum. The media was replaced by DMEM containing 0.1% calf serum for 24 h, after which the cells were stimulated with 15% calf serum and harvested at 0.5, 2, 4, 6 and 8 h. For harvesting, cells were washed twice with cold PBS, collected by scraping gently using a Cell Lifter (Corning Incorporated, 3008) and centrifuging at 1200 rpm for 3 min. Cells were harvested and used for phospho-proteomic analysis. Three independent replicate experiments were performed. 1×10^6^ A549 cells were seeded in 100mm plates and transduced with lentiviruses expressing control shRNA or three different TOLLIP shRNAs. Media was replaced the next day. After 24 h, cells were trypsinized, transferred to a 150mm plate and selected using puromycin (2.5µg/ml) for 2 days and harvested as described above. Three replicate experiments were performed for each shRNA construct.

### Phospho-proteomic analysis by mass spectrometry

Serum stimulated or unstimulated (0 h) NIH 3T3 cells were harvested (see above) and lysed in 500µl urea lysis buffer (8M Urea; 50mM Hepes pH 8.3), sonicated on ice 3 times at 1 min intervals followed by centrifugation at 14,000 rpm for 10 min at 4°C. Supernatants were collected and protein concentrations analyzed using BCA reagent (Bio-Rad). 250 mg protein was taken from each sample and the total volume was adjusted to 150 ul with 50 mM Hepes pH 8.3. The samples were then treated with 10mM DDT for 30 min at room temperature. Samples were alkylated using 20mM 2-Iodoacetamide for 30 min in the dark. 900µl cold acetone was added to each sample and left overnight at -20°C to precipitate proteins. The samples were centrifuged at 14,000 rpm for 10 min at 4°C and the protein precipitates were collected and resuspended in 50mM Hepes (pH 8.3). Next, the protein samples were treated with 6.5µg trypsin and incubated overnight at 37°C to generate tryptic peptides. The peptide samples were then dried under vacuum and resuspended in 100µl of 50mM Hepes (pH 8.3). Tandem mass tag (TMT) isobaric labeling (ThermoFisher, CA) was performed at 1:4 ratio by weight of sample for 1 h, and the reaction was quenched with 5% hydroxylamine. Different mass tags were used for peptides obtained from the NIH 3T3 cells harvested at each time point after serum stimulation; technical replicates received the same set of mass tags. Replicate experiments were analyzed in separate runs. TMT-labeled samples from each run were mixed, concentrated, dried under vacuum, and resuspended in 0.1% trifluoroacetic acid. To remove any uncoupled TMT, peptide samples were passed through empty C18 columns (ThermoFisher, CA). Eluted TMT labeled peptides were enriched for phosphopeptides by TiO_2_ affinity purification (High Select TiO_2_ kit; Thermo-Scientific, #A32993). The enriched phosphopeptides bound to TiO_2_ beads were eluted by high pH solvent and dried under vacuum. The flow through peptides and the phosphopeptides obtained from the various wash steps were collected separately, dried under vacuum, and resuspended in 0.1% TFA and further enriched for phosphopeptides using Fe-Oxide affinity purification (High Select Fe-NTA; Thermo-Scientific, #A32992). The phosphopeptides on the Fe-Oxide beads were then eluted by high pH solvents and combined with the TiO_2_ eluate collected earlier to obtain the maximum yield of phosphopeptides. The enriched phosphopeptides were fractionated by high pH (8.5) reverse phase (RP) on a Waters Acquity UPLC system coupled to a fluorescence detector (Waters, Milford, MA). The fractions were consolidated into 12 pools based on the chromatographic intensity. The flow-through and wash solutions of the Fe-oxide steps were collected, dried under vacuum, fractionated by high pH (8.5) RP chromatography, and consolidated into 24 pools based on the chromatographic intensity. The fraction pools were lyophilized and stored at -80°C until analysis by mass spectrometry.

For A549 cells, each TMT run contained four channels: an shCtrl sample and samples from three different TOLLIP shRNAs. The experiment was performed in triplicate, generating three replicates with four channels each. The replicate experiments were analyzed separately, resulting in 12 different phosphopeptide intensity measurements. TMT labeling and phosphopeptide enrichment were performed as described for NIH 3T3 cells. Different mass tags were used for the different cell types, while technical replicates received the same mass tags. Phosphopeptides were enriched using TiO_2_ affinity purification and Fe-Oxide affinity purification, fractionated and the resulting fraction pools were lyophilized and stored at -80°C.

### Mass spectrometry data acquisition and analysis

Data acquisition of TMT labeled peptides from NIH 3T3 cells was performed using Fusion Orbitrap (Thermo Scientific) and data for A549 cells were acquired using a Q-Exactive HF mass spectrometer (Thermo Scientific). The dried samples were reconstituted in 0.1% TFA and separated using a second dimension low pH gradient using a nanoflow liquid chromatography (Thermo Easy nLC 1000, Thermo Scientific) coupled to a mass spectrometer at 300 nl/min. The MS/MS analysis of the peptides along with the quantitative analysis of the TMT tags were performed at high resolution of 50,000 (Fusion)/45,000 (HF) in the orbitrap analyzer. Acquired MS/MS spectra were searched against mouse (for NIH 3T3) or human (for A549) Uniprot protein databases along with a contaminant protein database, using a SEQUEST and percolator validator algorithms in the Proteome Discoverer 2.2 software (Thermo Scientific, CA). The precursor ion tolerance was set at 10 ppm and the fragment ions tolerance was set at 0.02 Da along with methionine oxidation included as dynamic modification and TMT6 plex (229.163Da) set as a static modification of lysine and the N-termini of the peptide. For phospho-enriched peptides, phosphorylation of serine, threonine and tyrosine was set as a dynamic modification. Trypsin was specified as the proteolytic enzyme, with up to 2 missed cleavage sites allowed. A reverse decoy database search strategy was used to control for the false discovery rate and peptide identifications were validated using the percolator algorithm. Phosphopeptides and flow-through peptides from three independent experiments were analyzed. The affinity purified peptides were used to determine the abundance of phosphopeptides (denoted p-peptides), while the flow-through peptides were used to measure the protein abundance.

### Bioinformatic analysis of LC-MS data

All analyses using the MTM LC/MS/MS p-peptide and protein intensities were performed using either R (Version 4.3.1) or in-house programs. The twelve A549 samples (shCtrl, shTOLLIP.1, shTOLLIP.2 and shTOLLIP.3 groups, each transduced in triplicate) were analyzed using the mass tags described above. The first step was to consistently scale the intensities in these 12 channels for the p-peptides and proteins. Due to the varying number of missing intensities across the channels, a MAS5 scaling was found to produce the most consistent results. A p-peptide was removed from consideration if it did not have an observed intensity for at least two of the three technical replicates in each A549 group, if it did not represent a phosphorylated protein, or if the peptide sequence matched more than one protein. In certain cases, multiple p-peptides contain the same phosphorylated serine, threonine, or tyrosine residue. So that they were not treated independently, the Accession number and global position within the protein was used to sum the intensities at each phosphorylation site (p-site). The ratio of the p-site intensity to its corresponding protein intensity within the same channel was then determined. A p-site was removed if the p-site/protein ratio was not present in at least two of the three technical replicates for all cell cultures. The p-site/protein ratios were averaged across the two or three technical replicates for each group, followed by averaging the average ratios across the three TOLLIP knockdown groups. This overall TOLLIP knockdown p-site/protein ratio was divided by the average p-site/protein ratio for the shCtrl control samples to obtain a fold-change for each p-site. The log2-transform of these fold changes was converted to a z-scale and the p-sites that were below -1.5 were selected.

The residues surrounding each p-site were examined to determine potential matches to CK2, AKT or CDK phosphorylation motifs. A putative CK2-mediated p-site was identified if it passed one of the following two rules^42,66,67^, given that the p-site is at position *n*:

1. An aspartic or glutamic acid at either positions *n*+1 or *n*+3 or both, at least two acidic residues between positions *n*-1 and *n*+5, no proline or basic residues at position *n*+1, and no more than two basic residues between positions *n*-1 and *n*+5.
2. If there are no aspartic or glutamic acid at either positions *n*+1 or *n*+3, there must be at least three acidic residues between positions n-1 and n+6, no proline or basic residue at position *n*+1, and no more than two basic residues between positions *n*-1 and *n*+6.

A p-site was identified as a putative AKT target if an arginine was present at n-5 and n-3, and if a hydrophobic residue (alanine, phenylalanine, isoleucine, leucine, methionine, threonine, valine, or tryptophan) was present at n+1. Predicted CDK p-sites contained a proline at n+1 and a lysine or arginine at n+3.

The NIH 3T3 MTM data was treated similarly. Each of the three technical replicates at each time point was normalized using a MAS5 scaling. This was performed separately for the total protein and p-peptide data. Using the Accession number and global position within the protein, the p-peptide intensities were collapsed into p-site intensities. p-site/protein ratios were determined for each technical replicate at each time point. A p-site was removed if there were less than two replicates with valid p-site/protein ratios at any time point. The p-site/protein ratios were averaged for each time point. The average p-site/protein ratios at 0 min, 30 min and 2 h were then averaged. The average p-site/protein ratios at 4 or 6 h were then divided by the average of the average p-site/protein ratios at the 0-2 h time points, to represent the fold-change at 4 or 6 h. The log2-transformed fold-change across all p-sties was converted to a Z-score, and p-sites with a Z-score above 1.5 were considered up-regulated. For each set of up-regulated p-sites, the predicted CK2, AKT, and CDK p-sites were determined using the motif rules described above. Frequency distribution tables were generated using GraphPad prism 10.2.0. Using a bin width of 0.02, the number of values in each bin and the mean value per bin were plotted on the Y and X axes, respectively. Bin range was determined based on the distribution of the log2 fold changes within the selected phospho-proteomic data.

DAVID analysis^68^ was carried out for functional annotation of p-site lists; classifications based on Biological Processes (GOTERM_BP_DIRECT) were obtained and plotted. Ingenuity Pathway Analysis (IPA) software^69^ was used to perform “Phosphorylation Analysis” to generate phosphorylation/signaling networks. Proteins corresponding to the p-sites were used as identifiers and uploaded into the application. Networks generated by the IPA analysis are indicative of potential signaling cascades.

### RNA-sequencing

#### Library preparation

A total of 9 RNA samples (shCtrl, shTOLLIP.1 and shTOLLIP.2, each transduced in triplicate into A549 cells) were processed for library preparation using the Illumina Stranded Total RNA Prep and Ribo-Zero Plus. Prepared libraries were sequenced on an Illumina NovaSeq Xplus 10B platform. Paired-end reads were generated, with a total of 419 to 886 million reads per sample after filtering.

#### Data preprocessing and alignment

Raw sequencing data underwent initial quality control using FastQC^70^. Adapters and low-quality reads were removed using Cutadapt^71^. Clean reads were then aligned to the reference genome (hg38) using the STAR aligner^72^. In addition, the gene expression quantification analysis was performed for all samples using STAR/RSEM^73^.

#### Differential expression analysis

Differential expression analysis was performed using the DESeq2 package in R^74^. Raw counts were normalized and a negative binomial model was applied to identify genes with significantly altered expression between experimental conditions (shTOLLIP.1 and shTOLLIP.2 versus shCtrl, independently). Only genes differentially expressed in both conditions (shTOLLIP.1 versus shCtrl and shTOLLIP.2 versus shCtrl) were considered, based on an adjusted p-value cutoff of 0.05 and a mean fold change threshold of 0.5.

#### Gene set enrichment analysis (GSEA)

GSEA was conducted using the ClusterProfiler^75^ package in R. Pre-ranked gene lists, generated from the differential expression analysis results, were used as input for GSEA. Enrichment analysis was performed against gene sets obtained from the org.Hs.eg.db package^76^. The statistical significance of enrichment scores was assessed through permutation testing, and adjusted p-values were computed to account for multiple comparisons. The significance threshold for enriched gene sets was set at 0.05. Specific pathways analysis was conducted using the fgsea package^77^.

### Statistical analysis

Statistical analysis was performed using GraphPad Prism 10.2.0. Statistical significance for quantitation of immunoprecipitation experiments, RNPI, animal experiments (tumor lesion counts, tumor area, immunohistochemistry), colony formation assays, RNA-FISH and SA-βGal assays was calculated using two-tailed unpaired Student’s t test. P-values less than 0.05 were considered significant. The log-rank (Mantel-Cox) test was used for Kaplan–Meier survival analyses. Two-way ANOVA was used to analyze growth curves.

## References

1. Prior, I. A., Hood, F. E. & Hartley, J. L. The Frequency of Ras Mutations in Cancer. Cancer Research 80, 2969–2974 (2020).

2. Baines, A. T., Xu, D. & Der, C. J. Inhibition of Ras for cancer treatment: the search continues. Future Med Chem 3, 1787–1808 (2011).

3. Konieczkowski, D. J., Johannessen, C. M. & Garraway, L. A. A Convergence-Based Framework for Cancer Drug Resistance. Cancer Cell 33, 801–815 (2018).

4. Schneider, G., Schmidt-Supprian, M., Rad, R. & Saur, D. Tissue-specific tumorigenesis: context matters. Nat Rev Cancer 17, 239–253 (2017).

5. Tape, C. J., et al. Oncogenic KRAS Regulates Tumor Cell Signaling via Stromal Reciprocation. Cell 165, 1818 (2016).

6. Gober, M. K., Flight, R. M., Lambert, J., Moseley, H., Stromberg, A. & Black, E. P. Deregulation of a Network of mRNA and miRNA Genes Reveals That CK2 and MEK Inhibitors May Synergize to Induce Apoptosis KRAS-Active NSCLC. Cancer Inform 18, 1176935119843507 (2019).

7. Wang, H., et al. An integrative pharmacogenomics analysis identifies therapeutic targets in KRAS-mutant lung cancer. EBioMedicine 49, 106–117 (2019).

8. Papke, B. & Der, C. J. Drugging RAS: Know the enemy. Science 355, 1158–1163 (2017).

9. Moore, A. R., Rosenberg, S. C., McCormick, F. & Malek, S. RAS-targeted therapies: is the undruggable drugged? Nat Rev Drug Discov 19, 533–552 (2020).

10. The Cancer Genome Atlas Research Network. Comprehensive molecular profiling of lung adenocarcinoma. Nature 511, 543-550 (2014).

11. Yip-Schneider, M. T., Lin, A., Barnard, D., Sweeney, C. J. & Marshall, M. S. Lack of elevated MAP kinase (Erk) activity in pancreatic carcinomas despite oncogenic K-ras expression. Int J Oncol 15, 271–279 (1999).

12. Omerovic, J., Hammond, D. E., Clague, M. J. & Prior, I. A. Ras isoform abundance and signalling in human cancer cell lines. Oncogene 27, 2754–2762 (2008).

13. Yeh, J. J., et al. KRAS/BRAF mutation status and ERK1/2 activation as biomarkers for MEK1/2 inhibitor therapy in colorectal cancer. Mol Cancer Ther 8, 834–843 (2009).

14. Lim, K. H., et al. Activation of RalA is critical for Ras-induced tumorigenesis of human cells. Cancer Cell 7, 533–545 (2005).

15. Mizumoto, Y., et al. Activation of ERK1/2 occurs independently of KRAS or BRAF status in endometrial cancer and is associated with favorable prognosis. Cancer Sci 98, 652–658 (2007).

16. Murphy, J. E., Padilla, B. E., Hasdemir, B., Cottrell, G. S. & Bunnett, N. W. Endosomes: a legitimate platform for the signaling train. Proc Natl Acad Sci USA 106, 17615–17622 (2009).

17. Basu, S. K., et al. Oncogenic RAS-Induced Perinuclear Signaling Complexes Requiring KSR1 Regulate Signal Transmission to Downstream Targets. Cancer Research 78, 891–908 (2018).

18. Mellman, I. & Yarden, Y. Endocytosis and cancer. Cold Spring Harb Perspect Biol 5, a016949 (2013).

19. Sigismund, S., Confalonieri, S., Ciliberto, A., Polo, S., Scita, G. & Di Fiore, P. P. Endocytosis and signaling: cell logistics shape the eukaryotic cell plan. Physiol Rev 92, 273–366 (2012).

20. Jongsma, M. L., et al. An ER-Associated Pathway Defines Endosomal Architecture for Controlled Cargo Transport. Cell 166, 152–166 (2016).

21. Stewart, S., Sundaram, M., Zhang, Y., Lee, J., Han, M. & Guan, K. L. Kinase suppressor of Ras forms a multiprotein signaling complex and modulates MEK localization. Mol Cell Biol 19, 5523–5534 (1999).

22. Roy, F., Laberge, G., Douziech, M., Ferland-McCollough, D. & Therrien, M. KSR is a scaffold required for activation of the ERK/MAPK module. Genes Dev 16, 427–438 (2002).

23. Ritt, D. A., Zhou, M., Conrads, T. P., Veenstra, T. D., Copeland, T. D. & Morrison, D. K. CK2 Is a component of the KSR1 scaffold complex that contributes to Raf kinase activation. Curr Biol 17, 179–184 (2007).

24. Shtutman, M., et al. Function-based gene identification using enzymatically generated normalized shRNA library and massive parallel sequencing. Proc Natl Acad Sci USA 107, 7377–7382 (2010).

25. Wolf, J., et al. A mammosphere formation RNAi screen reveals that ATG4A promotes a breast cancer stem-like phenotype. Breast Cancer Res 15, R109 (2013).

26. Vizeacoumar, F. J., et al. A negative genetic interaction map in isogenic cancer cell lines reveals cancer cell vulnerabilities. Mol Syst Biol 9, 696 (2013).

27. Frodyma, D., Neilsen, B., Costanzo-Garvey, D., Fisher, K. & Lewis, R. Coordinating ERK signaling via the molecular scaffold Kinase Suppressor of Ras. F1000Res 6, 1621 (2017).

28. Michaud, N. R., et al. KSR stimulates Raf-1 activity in a kinase-independent manner. Proc Natl Acad Sci U S A 94, 12792–12796 (1997).

29. Jacobs, D., Glossip, D., Xing, H., Muslin, A. J. & Kornfeld, K. Multiple docking sites on substrate proteins form a modular system that mediates recognition by ERK MAP kinase. Genes Dev 13, 163–175 (1999).

30. Cacace, A. M., et al. Identification of constitutive and ras-inducible phosphorylation sites of KSR: implications for 14-3-3 binding, mitogen-activated protein kinase binding, and KSR overexpression. Mol Cell Biol 19, 229–240 (1999).

31. McKay, M. M., Ritt, D. A. & Morrison, D. K. Signaling dynamics of the KSR1 scaffold complex. Proc Natl Acad Sci USA 106, 11022–11027 (2009).

32. Ciarrocchi, A., et al. Tollip is a mediator of protein sumoylation. PLoS One 4, e4404 (2009).

33. Burns, K., et al. Tollip, a new component of the IL-1RI pathway, links IRAK to the IL-1 receptor. Nat Cell Biol 2, 346–351 (2000).

34. Shih, S. C., Prag, G., Francis, S. A., Sutanto, M. A., Hurley, J. H. & Hicke, L. A ubiquitin-binding motif required for intramolecular monoubiquitylation, the CUE domain. EMBO J 22, 1273–1281 (2003).

35. Yamakami, M., Yoshimori, T. & Yokosawa, H. Tom1, a VHS domain-containing protein, interacts with tollip, ubiquitin, and clathrin. J Biol Chem 278, 52865–52872 (2003).

36. Jumper, J., et al. Highly accurate protein structure prediction with AlphaFold. Nature 596, 583–589 (2021).

37. Sun, Y., Cheng, Z. & Liu, S. MCM2 in human cancer: functions, mechanisms, and clinical significance. Mol Med 28, 128 (2022).

38. Tsuji, T., Ficarro, S. B. & Jiang, W. Essential role of phosphorylation of MCM2 by Cdc7/Dbf4 in the initiation of DNA replication in mammalian cells. Mol Biol Cell 17, 4459–4472 (2006).

39. Li, X., Goobie, G. C. & Zhang, Y. Toll-interacting protein impacts on inflammation, autophagy, and vacuole trafficking in human disease. Journal of molecular medicine 99, 21–31 (2021).

40. Begka, C., et al. Toll-Interacting Protein Regulates Immune Cell Infiltration and Promotes Colitis-Associated Cancer. iScience 23, 100891 (2020).

41. Purzner, T., et al. Developmental phosphoproteomics identifies the kinase CK2 as a driver of Hedgehog signaling and a therapeutic target in medulloblastoma. Sci Signal 11, (2018).

42. Meggio, F. & Pinna, L. A. One-thousand-and-one substrates of protein kinase CK2? FASEB J 17, 349–368 (2003).

43. Widmann, B., et al. The kinase activity of human Rio1 is required for final steps of cytoplasmic maturation of 40S subunits. Mol Biol Cell 23, 22–35 (2012).

44. Turowski, T. W., et al. Rio1 mediates ATP-dependent final maturation of 40S ribosomal subunits. Nucleic Acids Res 42, 12189–12199 (2014).

45. Plassart, L., et al. The final step of 40S ribosomal subunit maturation is controlled by a dual key lock. eLife 10, (2021).

46. Ameismeier, M., et al. Structural basis for the final steps of human 40S ribosome maturation. Nature 587, 683–687 (2020).

47. Luo, J., et al. A genome-wide RNAi screen identifies multiple synthetic lethal interactions with the Ras oncogene. Cell 137, 835–848 (2009).

48. Weinberg, F., et al. The Atypical Kinase RIOK1 Promotes Tumor Growth and Invasive Behavior. EBioMedicine 20, 79–97 (2017).

49. Hong, X., et al. Targeting posttranslational modifications of RIOK1 inhibits the progression of colorectal and gastric cancers. eLife 7, (2018).

50. Kubinski, K. & Maslyk, M. The Link between Protein Kinase CK2 and Atypical Kinase Rio1. Pharmaceuticals 10, (2017).

51. Toruń, A., et al. Endocytic Adaptor Protein Tollip Inhibits Canonical Wnt Signaling. PLoS One 10, e0130818 (2015).

52. Zhu, L., et al. Tollip, an intracellular trafficking protein, is a novel modulator of the transforming growth factor-β signaling pathway. J Biol Chem 287, 39653–39663 (2012).

53. Didierlaurent, A., et al. Tollip regulates proinflammatory responses to interleukin-1 and lipopolysaccharide. Mol Cell Biol 26, 735–742 (2006).

54. Pokatayev, V., et al. Homeostatic regulation of STING protein at the resting state by stabilizer TOLLIP. Nat Immunol 21, 158–167 (2020).

55. Thapa, N., Chen, M., Horn, H. T., Choi, S., Wen, T. & Anderson, R. A. Phosphatidylinositol-3-OH kinase signalling is spatially organized at endosomal compartments by microtubule-associated protein 4. Nat Cell Biol 22, 1357–1370 (2020).

56. Pelletier, J., Thomas, G. & Volarevic, S. Ribosome biogenesis in cancer: new players and therapeutic avenues. Nat Rev Cancer 18, 51–63 (2018).

57. Azman, M. S., et al. An ERK1/2-driven RNA-binding switch in nucleolin drives ribosome biogenesis and pancreatic tumorigenesis downstream of RAS oncogene. EMBO J 42, e110902 (2023).

58. Feldser, D. M., et al. Stage-specific sensitivity to p53 restoration during lung cancer progression. Nature 468, 572–575 (2010).

59. Johnson, L., et al. Somatic activation of the K-ras oncogene causes early onset lung cancer in mice. Nature 410, 1111–1116 (2001).

60. Tuveson, D. A., et al. Endogenous oncogenic K-ras(G12D) stimulates proliferation and widespread neoplastic and developmental defects. Cancer Cell 5, 375–387 (2004).

61. Hingorani, S. R., et al. Trp53R172H and KrasG12D cooperate to promote chromosomal instability and widely metastatic pancreatic ductal adenocarcinoma in mice. Cancer Cell 7, 469–483 (2005).

62. Therrien, M., Michaud, N. R., Rubin, G. M. & Morrison, D. K. KSR modulates signal propagation within the MAPK cascade. Genes Dev 10, 2684–2695 (1996).

63. Campeau, E., et al. A versatile viral system for expression and depletion of proteins in mammalian cells. PLoS One 4, e6529 (2009).

64. Boothe, T., et al. Inter-domain tagging implicates caveolin-1 in insulin receptor trafficking and Erk signaling bias in pancreatic beta-cells. Mol Metab 5, 366–378 (2016).

65. Schindelin, J., et al. Fiji: an open-source platform for biological-image analysis. Nat Methods 9, 676-682 (2012).

66. St-Denis, N., et al. Systematic investigation of hierarchical phosphorylation by protein kinase CK2. J Proteomics 118, 49–62 (2015).

67. Wang, C., et al. Determination of CK2 specificity and substrates by proteome-derived peptide libraries. Journal of proteome research 12, 3813–3821 (2013).

68. Sherman, B. T., et al. DAVID: a web server for functional enrichment analysis and functional annotation of gene lists (2021 update). Nucleic Acids Res, (2022).

69. Kramer, A., Green, J., Pollard, J., Jr. & Tugendreich, S. Causal analysis approaches in Ingenuity Pathway Analysis. Bioinformatics 30, 523–530 (2014).

70. Babraham Bioinformatics. A quality control tool for high throughput sequence data. (ed(eds). Babraham Institute (2023).

71. Martin, M. Cutadapt removes adapter sequences from high-throughput sequencing reads. In: EMBnet (ed(eds) (2011).

72. Dobin, A., et al. STAR: ultrafast universal RNA-seq aligner. Bioinformatics 29, 15–21 (2013).

73. Li, B. & Dewey, C. N. RSEM: accurate transcript quantification from RNA-Seq data with or without a reference genome. BMC Bioinformatics 12, 323 (2011).

74. Love, M. I., Huber, W. & Anders, S. Moderated estimation of fold change and dispersion for RNA-seq data with DESeq2. Genome Biol 15, 550 (2014).

75. Wu, T., et al. clusterProfiler 4.0: A universal enrichment tool for interpreting omics data. Innovation (Camb) 2, 100141 (2021).

76. Carlson, M. org.Hs.eg.db: Genome wide annotation for Human. (ed(eds). Bioconductor (2022).

77. Korotkevich, G., Sukhov, V., Budin, N., Shpak, B., Artyomov, M. N. & Sergushichev, A. Fast gene set enrichment analysis. bioRxiv, 060012 (2021).

