## Supplementary Figures and legends for "CK2 signaling from TOLLIP-dependent perinuclear endosomes is an essential feature of *KRAS* and *NRAS* mutant cancers"

### Supplementary Figure 1

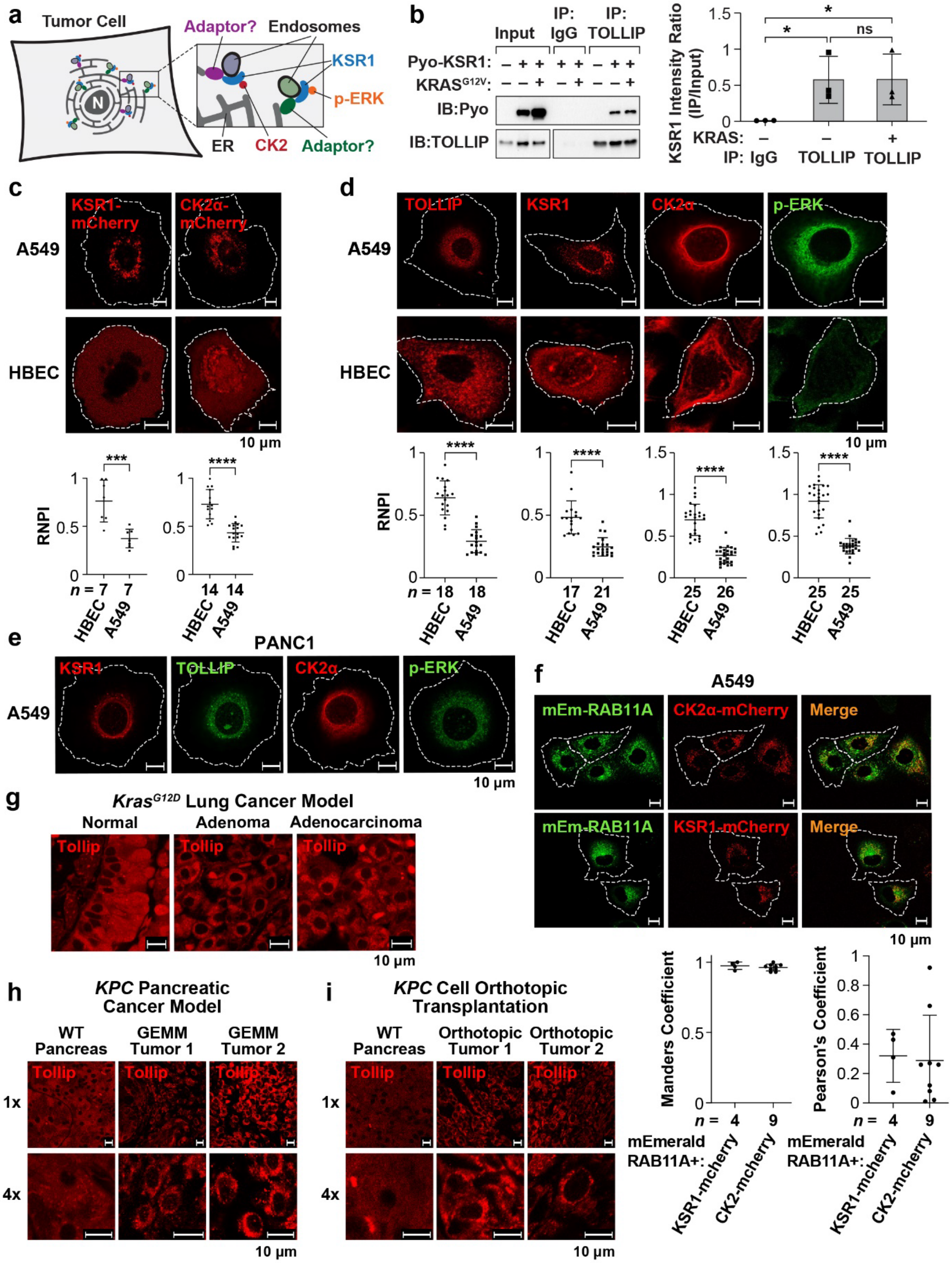

**Supplementary Fig. 1 | Perinuclear TOLLIP corresponds to the presence of PSCs in tumor cell lines and tissues. a** A model for recruitment of signaling endosomes to the perinuclear ER network by adaptor proteins. CK2 and ERK are present on different types of signaling endosomes that use adaptor proteins for attachment to the ER. **b** IP experiment showing endogenous TOLLIP associating with ectopically expressed Pyo-KSR1, without or with $KRAS^{G12V}$, in 293T cells. co-IP data (right) are mean ± s.d.; $n = 3$ independent experiments. **c** Fluorescently-tagged KSR1 and CK2a are perinuclear in A549 cells and pan-cytoplasmic in HBEC cells. Bottom: RNPI data showing nuclear proximity of each protein in the two cell lines; *n*, number of cells analyzed. **d** IF imaging illustrating perinuclear clustering of TOLLIP, KSR1, CK2a and p-ERK in A549 cells and dispersed cytoplasmic distribution in non-transformed HBEC cells. Bottom: RNPI data showing nuclear proximity of each protein in the two cell lines; *n*, number of cells analyzed. **e** Endogenous TOLLIP, CK2a and p-ERK segregate to the nuclear-proximal region in *KRAS* mutant PANC1 (PDAC) cells. **f** Live cell imaging showing partial co-localization of fluorescently-tagged RAB11A with CK2α and KSR1 in A549 cells. Bottom: RAB11A colocalization with CK2α and KSR1 was quantified using Manders *r* and Pearson's *R*; *n*, number of cells analyzed. **g** Tollip is perinuclear in $Kras^{G12D}$-driven mouse lung adenomas and ADCs but shows diffuse immunostaining in unaffected normal lung tissue. **h** Tollip forms perinuclear rings in primary PDAC tumors arising in *KPC* mice, compared to pan-cytoplasmic distribution in normal pancreas. **i** Tollip displays perinuclear staining in pancreatic tumors from orthotopic *KPC* cell allografts versus uniform distribution in normal host pancreas.

Statistical significance for co-IP assays and RNPI data were determined using two-tailed unpaired Student's t test; $^{*}P \leq 0.05$, $^{***}P \leq 0.001$, $^{****}P \leq 0.0001$.

### Supplementary Figure 2

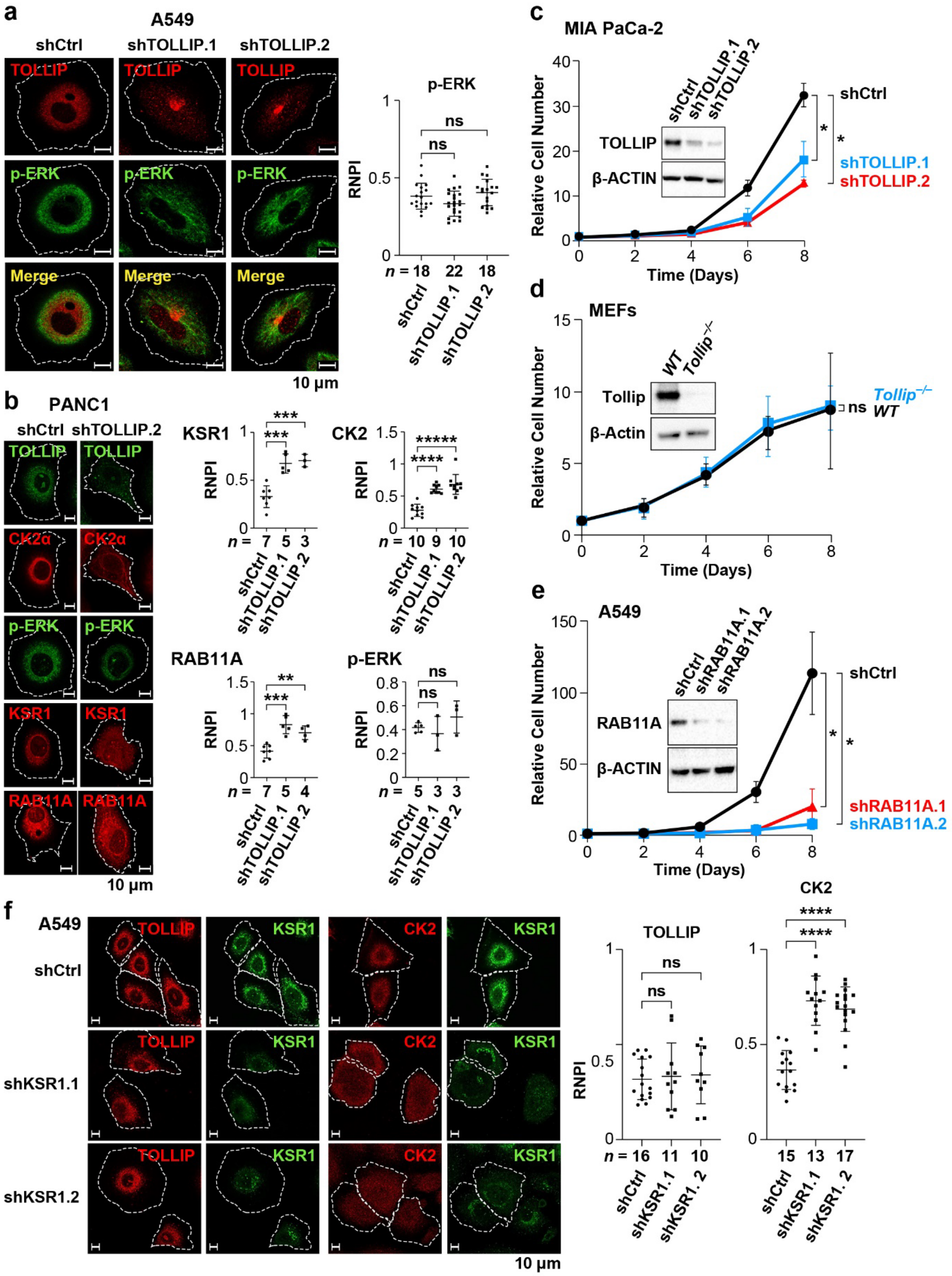

**Supplementary Fig. 2 | Roles of TOLLIP, RAB11A and KSR1 in PSC formation. a** TOLLIP depletion in A549 cells does not alter perinuclear localization of p-ERK. Right: RNPI analysis of p-ERK nuclear proximity; *n*, number of cells analyzed. **b** TOLLIP silencing disrupts perinuclear localization of CK2α, KSR1 and RAB11A but not p-ERK in PANC1 cells ($KRAS^{G12C}$ PDAC). Right: RNPI analysis of each protein in control and TOLLIP-depleted cells; *n*, number of cells analyzed. **c** TOLLIP knockdown decreases proliferation of MIA PaCa-2 cells ($KRAS^{G12C}$ PDAC). Data are mean ± s.d.; $n$ = 3 independent experiments. **d** *WT* and $Tollip^{-/-}$ MEFs proliferate at similar rates. Data are mean ± s.d.; $n$ = 3 independent experiments. **e** RAB11A silencing impairs proliferation of A549 cells. Data are mean ± s.d.; $n$ = 3 independent experiments. **f** A549 cells depleted for KSR1 retain perinuclear localization of TOLLIP but not CK2α. Right: RNPI analysis of each protein in control and KSR1-depleted cells; *n*, number of cells analyzed.

Two-way ANOVA was used to analyze growth curves. RNPI data was analyzed using two-tailed unpaired Student's *t*-test; $*P \leq 0.05$, $**P \leq 0.01$, $***P \leq 0.001$, $****P \leq 0.0001$.

### Supplementary Figure 3

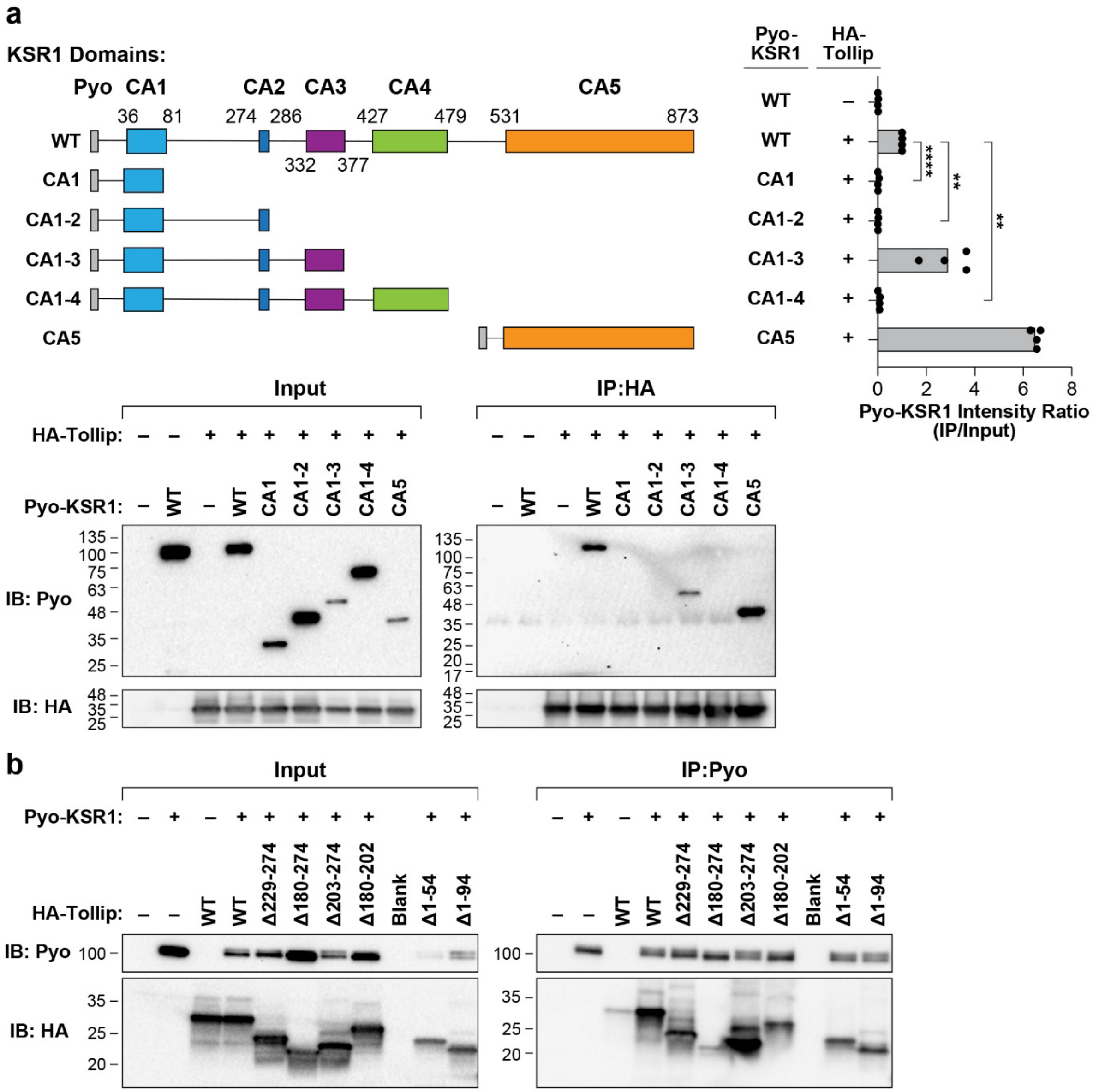


**c**

```
FRUITFLY – D. lebanonensis     176 EGSIDIVLSYSLQPFTYPIAGTS-------FLVGSNARPIVLTP-------VHQA----- 216
SPIDER – A. ventricosus        175 EGTINLVITCQPIPSGNLIYHPK------SLVTIVPFGPFSDEEGVVSDRPVSHVNDQSR 228
NEMATODE – C. elegans          162 EGMIHLHFSFAPIDLPLQQA---------------------------------------- 181
FROG – X. laevis               139 EGMINLVMSYTSVPA--MMPAQPVVLMPTVYQQGVGYVPIAD--PIYNPGMP-------- 210
ZEBRAFISH – D. rerio           163 EGMINLVMSFATIPAGMMMQPQPVVLMPTVYQQGVGYVPIAGVPGMYNQGVVPMG----- 193
HUMAN – H. sapiens             163 EGMINLVMSYALLPAAMVMPPQPVVLMPTVYQQGVGYVPITGMPAVCSPGMVPVA----- 217
RAT – R. norvegicus            163 EGMINLVMSYTSLPAAMMMPPQPVVLMPTVYQQGVGYVPIAGMPAVCSPGMVPMA----- 217
MOUSE – M. musculus            163 EGMINLVMSYTSLPAAMMMPPQPVVLMPTVYQQGVGYVPIAGMPAVCSPGMVPMA----- 217
RHESUS-MONKEY – M. mulatta     163 EGMINLVMSYTSLPAAMMMPPQPVVLMPTVYQQGVGYVPIAGMPAVCSPGMVPMA----- 217
                                   ** *.: ::              185                202
```

**Supplementary Fig. 3 | Mapping the interaction domains in KSR1 and Tollip. a** The conserved KSR1 CA5 pseudokinase domain mediates association with Tollip. Nested KSR1 C-terminal deletions were generated to map the Tollip binding region. Conserved regions CA1-CA5 and the endpoint of each deletion are depicted, with a Pyo tag appended to the N termini. The CA5 domain alone was also tested. KSR1 proteins were co-expressed with HA-Tollip in HEK293T cells and lysates were immunoprecipitated using an HA antibody. Bottom: input lysates and IP samples were analyzed by immunoblotting for HA or Pyo. A representative blot out of four independent experiments is shown. Upper right: band intensities were quantified and IP/input ratios determined for each KSR1 protein; values were normalized to WT KSR1. Data are means ± s.d.; $n$ = 4 independent experiments. **b** Immunoblot showing HA-Tollip mutants co-immunoprecipitating with Pyo-KSR1 (see Fig. 3a). A representative blot out of three independent experiments is shown. **c** The KSR1 binding region of Tollip includes an 18 aa core sequence (185-202) that is conserved among vertebrate species. The alignment shows six vertebrate and three invertebrate Tollip sequences surrounding the core KSR1 binding motif (highlighted).

co-IP data was analyzed using two-tailed unpaired Student's $t$-test; $^{**}P \leq 0.01$, $^{****}P \leq 0.0001$.

Supplementary Figure 4

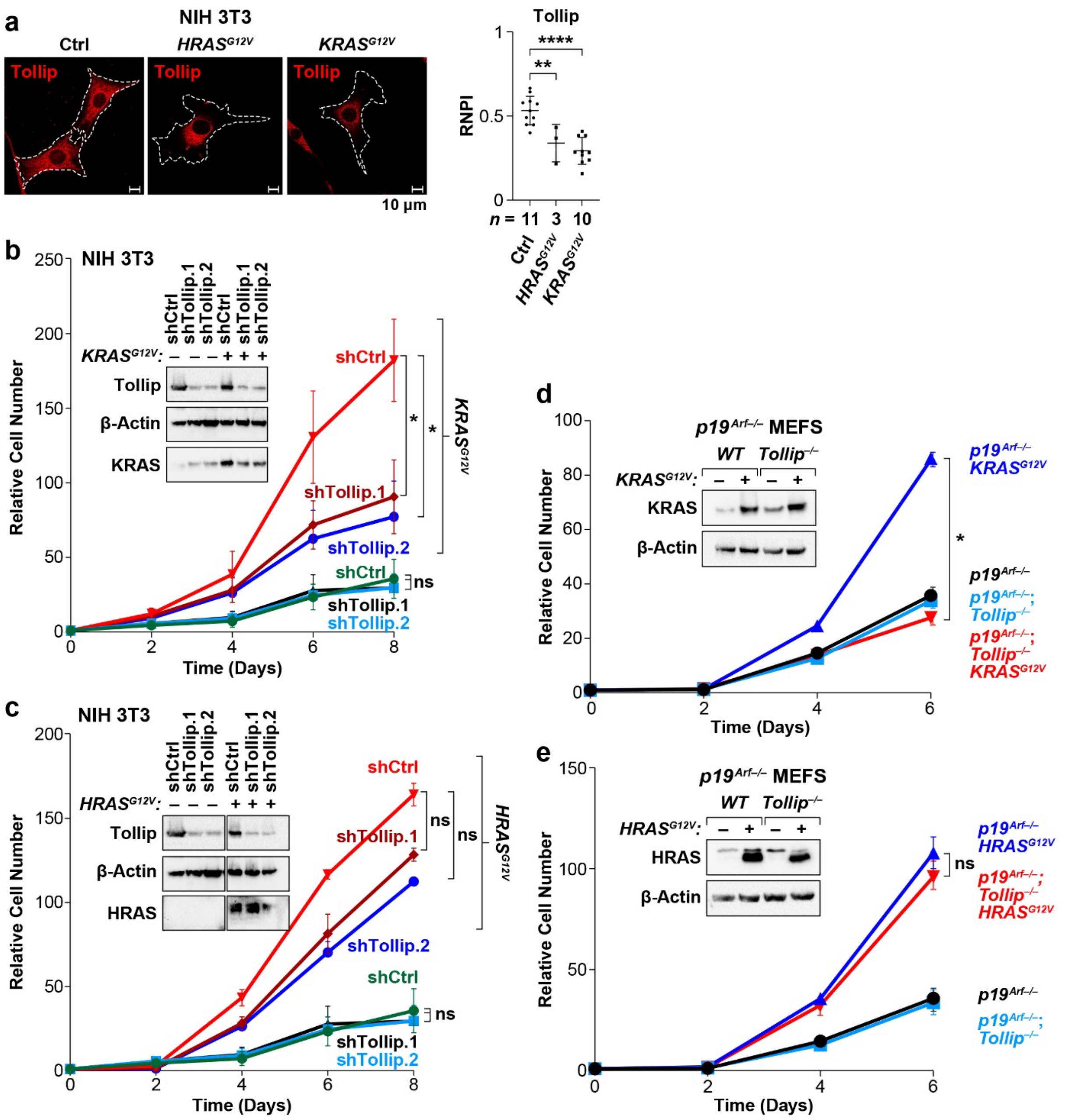


**Supplementary Fig. 4 | TOLLIP dependence of various transformed/tumor cell lines. a** Oncogenic *KRAS* and *HRAS* induce perinuclear localization of Tollip in NIH3T3 cells. Right: RNPI analysis of Tollip nuclear proximity in each cell population; *n*, number of cells analyzed. **b**,**c** Proliferation of control or *RAS*-transformed NIH 3T3 cells without and with Tollip depletion. **b** $KRAS^{G12V}$ cells; **c** $HRAS^{G12V}$ cells. The same growth curves for non-transformed control cells are shown in the two figures (a single experiment was performed using common control cells, and the data for *KRAS*$^{G12V}$ and *HRAS*$^{G12V}$ were separated for clarity). Data are mean ± s.d.; *n* = 3 independent experiments. **d,e** *p19*$^{Arf-/-}$ MEFs require Tollip for *KRAS*$^{G12V}$- but not *HRAS*$^{G12V}$-driven hyperproliferation. **d** *KRAS*$^{G12V}$ cells; **e** *HRAS*$^{G12V}$ cells. *WT* and *Tollip*$^{-/-}$ MEFs were nullizygous for *p19*$^{Arf}$ to allow transformation by oncogenic RAS. Data are mean ± s.d.; *n* = 3 independent experiments.

Two-way ANOVA was used to analyze growth curves and two-tailed unpaired Student's t-test was used to evaluate RNPI data; *$P \leq 0.05$, **$P \leq 0.01$, ****$P \leq 0.0001$.

### Supplementary Figure 5

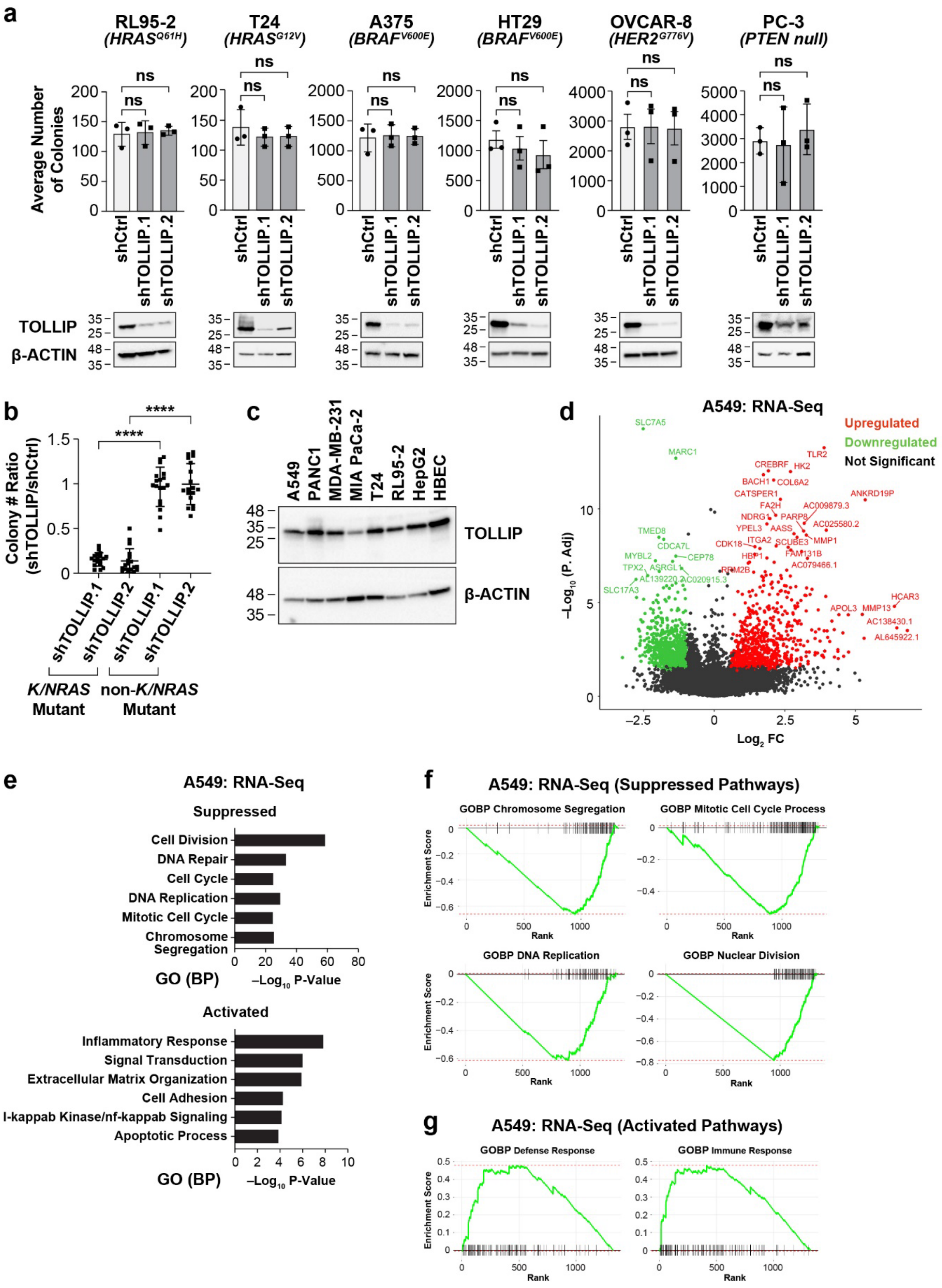

**Supplementary Fig. 5 | Characterization of TOLLIP dependence in human tumor cell lines and a TOLLIP-dependent transcriptome in A549 cells. a** Human tumor cells carrying oncogenic mutations in *HRAS* (RL95-2, endometrial carcinoma; T24, bladder carcinoma), *BRAF* (A375, melanoma; HT29, colorectal adenocarcinoma), *ERBB2* (OVKAR-8, ovarian carcinoma) or *PTEN* (PC3, prostatic small cell carcinoma) are unaffected by TOLLIP depletion in clonogenic growth assays. Data are mean ± s.d.; $n$ = 3 independent experiments. **b** Amalgamated data comparing the TOLLIP dependencies of *K/NRAS* mutant and non-*K/NRAS* mutant tumor cells. Clonogenic growth data from **a** and Fig. 4d were used to generate shTOLLIP/shCrtl ratios for the three biological replicates from each tumor cell line; the combined results are separated according to *K/NRAS* mutant status. **c** TOLLIP dependency of tumor cell lines does not correlate with its expression levels. Several of the cell lines tested for TOLLIP dependence in clonogenic assays were analyzed by immunoblotting for TOLLIP; β-ACTIN, loading control. **d** Volcano plot of RNA-seq data showing genes whose expression changes in TOLLIP-depleted A549 cells. FC, fold change (shTOLLIP/shCtrl). **e** GO categories (biological process) for mRNAs that are up- or down-regulated following TOLLIP knockdown. **f**, Gene set enrichment analysis (GSEA) of down-regulated mRNAs in TOLLIP-depleted cells. Four of the top pathways are shown. **g** Gene set enrichment analysis (GSEA) of up-regulated mRNAs in TOLLIP-depleted cells. Two of the top pathways are shown.

Two-tailed unpaired Student's t-test was performed to evaluate clonogenic growth data; $****P \leq 0.0001$.

Supplementary Figure 6

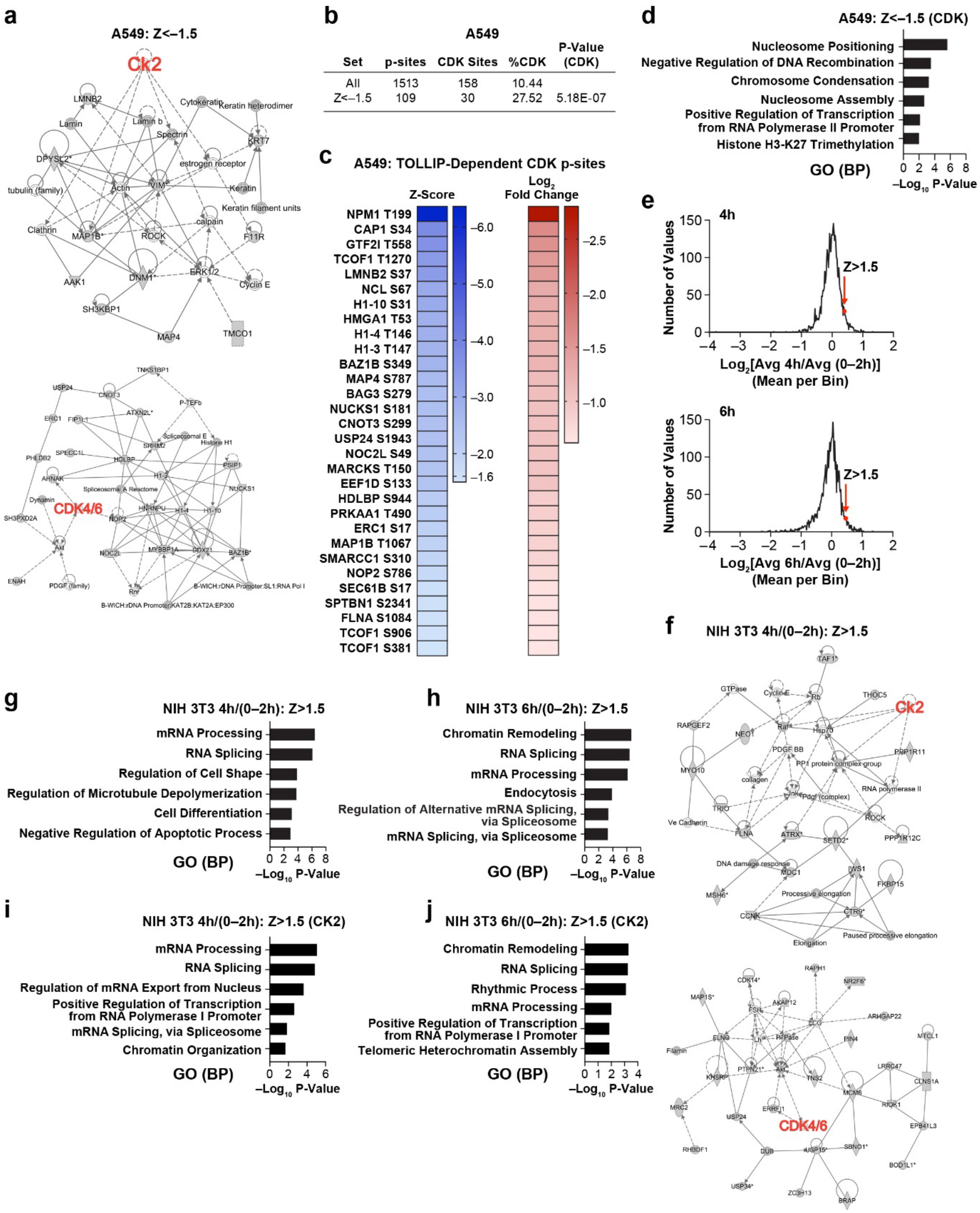


**Supplementary Fig. 6 | Bioinformatic analysis of phosphoproteomic data from A549 cells and NIH 3T3 cells. a** Signaling networks (Ingenuity Pathway Analysis; IPA) corresponding to p-sites significantly reduced ($Z \leq -1.5$) in TOLLIP-depleted A549 cells. Two of the three highest-scoring networks are shown; CK2 and CDK nodes are indicated in red. **b** Enrichment of predicted CDK motifs in p-sites that decrease ($Z \leq -1.5$) in TOLLIP-depleted A549 cells. Comparison is between all p-sites and those that are down-regulated upon TOLLIP knockdown. **c** List of predicted CDK sites from the set of down-regulated p-sites ($Z \leq -1.5$) in A549 cells. Heat maps depict Z-score and $\log_2$ fold change for each site. **d** Ranked GO categories (biological process) for the set of down-regulated CDK-like p-sites shown in panel **c**. **e** Frequency distribution curves for p-sites at 4 h and 6 h divided by the combined average intensity of the 0-2 h time points in serum-stimulated NIH 3T3 cells. $Z > 1.5$ cutoffs are indicated. **f** IPA signaling networks corresponding to p-sites increased at 4 h in serum-stimulated NIH 3T3 cells ($Z \geq 1.5$). Two of the four highest-scoring networks are shown; CK2 and CDK nodes are indicated in red. **g**,**h** Ranked GO categories for all p-sites up-regulated ($Z \geq 1.5$) in serum-stimulated NIH 3T3 cells at 4 h (**g**) or 6 h (**h**). **i**,**j** Ranked GO categories for predicted CK2 p-sites up-regulated ($Z \geq 1.5$) in serum-stimulated NIH 3T3 cells at 4 h (**i**) or 6 h (**j**).

Statistical significance of CDK site enrichment in the $Z < -1.5$ p-sites compared to all p-sites (**b**) was assessed using the one-tailed Binomial test.

Supplementary Figure 7

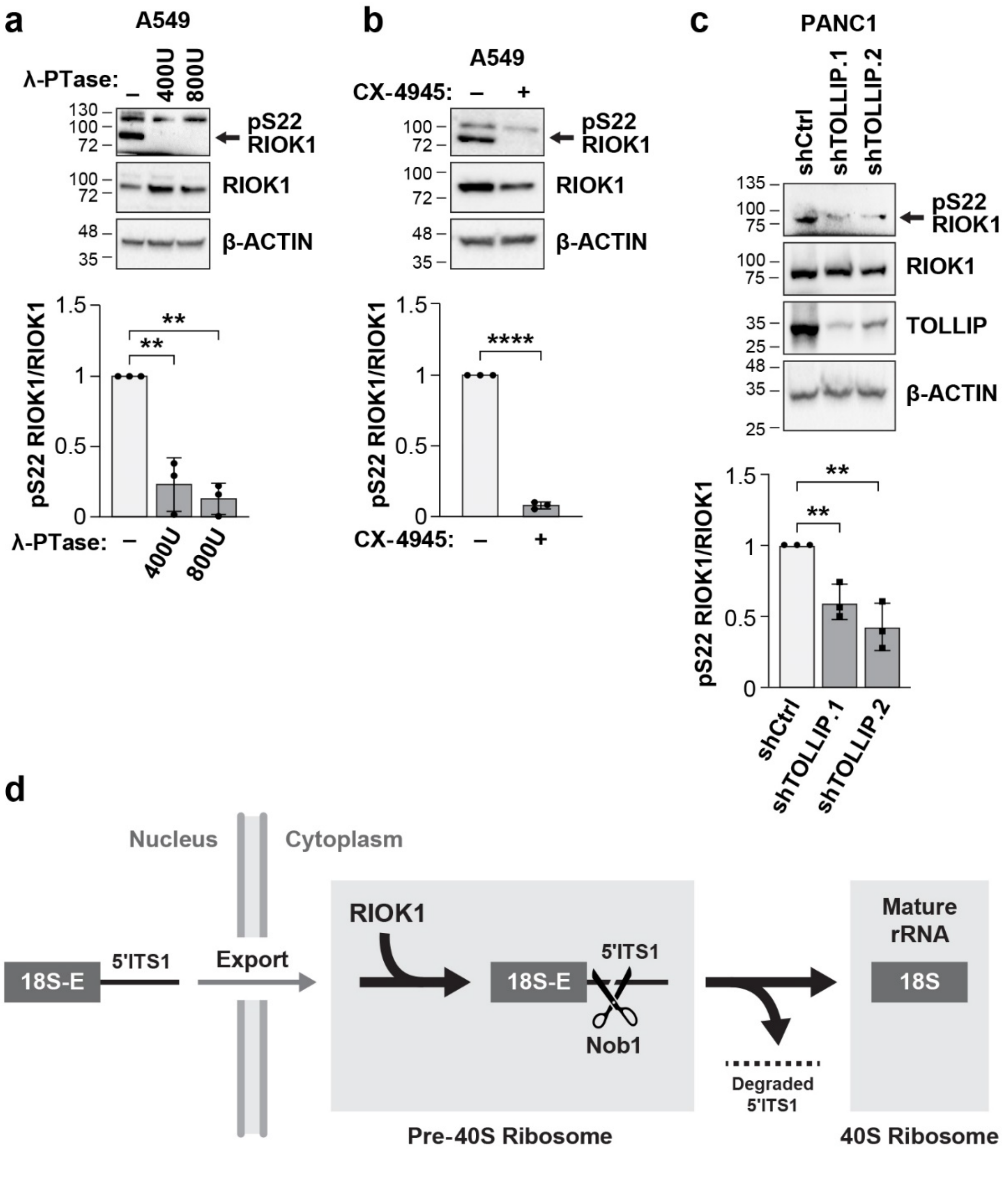


**Supplementary Fig. 7 | Analysis of RIOK1 Ser22 phosphorylation and its dependence on TOLLIP. a** λ-phosphatase treatment decreases pSer22 RIOK1 detection in lysates from A549 cells. pSer22 RIOK1 immunoblot data normalized to total RIOK1 (bottom) are mean ± s.d.; *n* = 3 independent experiments. **b** Treatment of A549 cells with the CK2 inhibitor CX-4945 decreases pSer22 RIOK1 levels. Immunoblot data normalized to total RIOK1 (bottom) are mean ± s.d.; *n* = 3 independent experiments. **c** TOLLIP knockdown diminishes RIOK1 Ser22 phosphorylation in

PANC1 cells. pSer22 RIOK1 levels normalized to total RIOK1 (bottom) are mean ± s.d.; $n = 3$ independent experiments. **d** Diagram showing the role of RIOK1 in final steps of 18S rRNA processing and 40S ribosome maturation. In the absence of RIOK1, the 18S-E precursor is not cleaved by Nob1 and the 5'ITS1 spacer sequence remains associated with the pre-40S ribosome in the cytoplasm and is protected from degradation. See the main text for references.

Two-tailed unpaired Student's t-test was used to evaluate changes in pSer22 RIOK1 levels; $**P \leq 0.01$, $****P \leq 0.0001$.

### Supplementary Figure 8

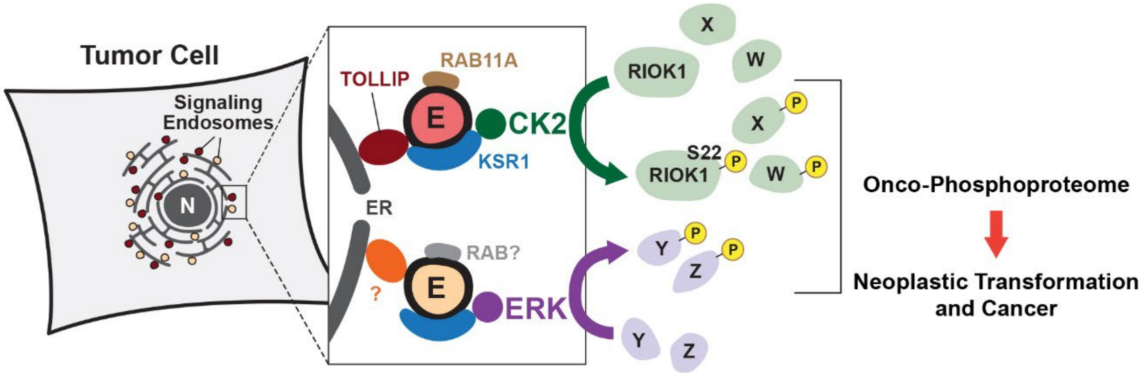


**Supplementary Fig. 8 | Summary diagram.** Model showing perinuclear ER tethering of CK2 and ERK1/2 signaling endosomes via endosomal adapter proteins in tumor cells. *K/NRAS* mutant cells use TOLLIP as the CK2 endosomal adapter, while a distinct but functionally related adapter is presumably employed in non-*K/NRAS* mutant cells. ERK endosomes are tethered by a different adapter protein, yet to be identified. Both CK2 and ERK1/2 are recruited to endosomes through KSR1. These and presumably other perinuclear kinases access a selective set of substrates, including RIOK1 (CK2 target), generating an onco-phosphoproteome that drives neoplastic transformation and tumorigenesis.
